# Overexpression of key sterol pathway enzymes in two model marine diatoms alters sterol profiles in *Phaeodactylum tricornutum*

**DOI:** 10.1101/2020.07.30.228171

**Authors:** Ana Cristina Jaramillo-Madrid, Raffaela Abbriano, Justin Ashworth, Michele Fabris, Peter J. Ralph

## Abstract

Sterols are a class of triterpenoid molecules with diverse functional roles in eukaryotic cells, including intracellular signaling and regulation of cell membrane fluidity. Diatoms are a dominant eukaryotic phytoplankton group that produce a wide diversity of sterol compounds. The enzymes 3-hydroxy-3-methyl glutaryl CoA reductase (*HMGR*) and squalene epoxidase (SQE) have been reported to be rate-limiting steps in sterol biosynthesis in other model eukaryotes; however, the extent to which these enzymes regulate triterpenoid production in diatoms is not known. To probe the role of these two metabolic nodes in the regulation of sterol metabolic flux in diatoms, we independently over-expressed two versions of the native *HMGR* and a conventional, heterologous SQE gene in the diatoms *Thalassiosira pseudonana* and *Phaeodactylum tricornutum*. Overexpression of these key enzymes resulted in significant differential accumulation of downstream sterol pathway intermediates in *P. tricornutum*. HMGR-mVenus overexpression resulted in the accumulation of squalene, cycloartenol, and obtusifoliol, while cycloartenol and obtusifoliol accumulated in response to heterologous NoSQE-mVenus overexpression. In addition, accumulation of the end-point sterol 24-methylenecholesta-5,24(24’)-dien-3β-ol was observed in all *P. tricornutum* overexpression lines, and campesterol increased 3-fold in *P. tricornutum* lines expressing NoSQE-mVenus. Minor differences in end-point sterol composition were also found in *T. pseudonana*, but no accumulation of sterol pathway intermediates was observed. Despite the successful manipulation of pathway intermediates and individual sterols in *P. tricornutum*, total sterol levels did not change significantly in transformed lines, suggesting the existence of tight pathway regulation to maintain total sterol content.

## INTRODUCTION

Sterols are essential triterpenoids that function as regulators of cell membrane dynamics in all eukaryotic organisms (Dufourc, 2008). In animals and higher plants, sterols participate in the synthesis of secondary metabolites involved in defense mechanisms, and steroid hormones that regulate growth and development (Valitova *et al*., 2016). Due to their presence in ancient sediments, sterol compounds are used as durable biomarkers to track important evolutionary events (Gold *et al*., 2017). Sterols of plant origin, known as phytosterols, are used as nutraceuticals for their cholesterol-lowering effects (Ras *et al*., 2014). Other therapeutic applications such as anti-inflammatory (Aldini *et al*., 2014) and anti-diabetic activities (Wang *et al*., 2017) are currently under research. In order to meet increasing demands in the global phytosterols market, about 7–9% per annum (Borowitzka, 2013), diatoms have been proposed as an alternative source of sterols (Jaramillo-Madrid *et al*., 2019).

Diatoms are primary constituents of phytoplankton communities and principal players in the global carbon cycle. These photosynthetic microorganisms are an important ecological group of microalgae present in a great diversity of aquatic environments (Armbrust, 2009). Diatoms exhibit higher photosynthetic efficiencies than plants and are adaptable to environmental challenges encountered in dynamic and competitive marine environments (Hildebrand *et al*., 2012), which are also characteristics suited to the microbial production of bioproducts. Diatoms are emerging as alternative and sustainable hosts for terpenoids production (D’Adamo *et al*., 2019; Fabris *et al*., 2020). As complex organisms with a particular evolution history, diatoms possess a unique metabolism (Fabris *et al*., 2012, 2014; Pollier *et al*., 2019; Jaramillo-Madrid *et al*., 2020a) that can represent an advantage for production of terpenoid such as sterols (Vavitsas *et al*., 2018).

Diatoms produce high proportions of a large variety of sterol compounds (Rampen *et al*., 2010). Sterol sulfates appear to be important regulators of diatom bloom dynamics, as they were shown to trigger programmed cell death in the marine diatom *Skeletonema marinoi* (Gallo *et al*., 2017). Recent studies suggest that sterol biosynthesis is tightly regulated. Levels of intermediate compounds in sterol synthesis change in response to different environmental conditions (Jaramillo-Madrid *et al*., 2020b) and to the addition of chemical inhibitors (Jaramillo-Madrid *et al*., 2020a). However, end-point sterol levels remained unchanged under the same treatments. Deeper understanding of diatom sterol metabolism will provide ecological insights as well as enable future metabolic engineering efforts for biotechnological applications. In particular, the regulation of the sterol biosynthesis in diatoms is not yet well understood.

Isoprenoid sterol precursors can be synthesized through either the cytosolic mevalonate (MVA) pathway or the plastidial methylerithriol phosphate (MEP) pathway. In most eukaryotic organisms, only one of the two pathways are present (Lohr *et al*., 2012). However, in plants, both pathways are functional but the MVA provides the substrates for sterol biosynthesis (Vranova *et al*., 2013). In diatoms, including the model diatoms *Thalassiosira pseudonana* and *Phaeodactylum tricornutum*, both pathways are functional (Jaramillo-Madrid *et al*., 2020a). However, there is no evidence for MVA presence in some diatoms such as *Haslea ostrearia* and *Chaetoceros muelleri* (Massé *et al*., 2004; Athanasakoglou *et al*., 2019; Jaramillo-Madrid *et al*., 2020a). In these diatoms, synthesis of isoprenoids may rely solely on the MEP pathway, as is the case for some green and red algae (Lohr et al., 2012). The presence of both the MVA and MEP pathways is an advantage for engineering efforts, as it potentially provides a higher pool of intermediates for isoprenoid production (Sasso *et al*., 2012; Jaramillo-Madrid *et al*., 2020a). It has been recently demonstrated that in *P. tricornutum* products from MVA pathway accumulated in the cytoplasm can be used for the production of non-endogenous terpenoids such as geraniol, indicating presence of free GPP pool (Fabris *et al*., 2020).

In the MVA pathway, three molecules of acetyl-CoA are transformed into isopentenyl diphosphate (IPP) and dimethylallyl diphosphate (DMAPP) (Fig. 1). In plants, fungi, and animals, the 3-hydroxy-3-methylglutaryl-co-enzyme-A reductase (*HMGR*, E.C. 1.1.1.34) is one of the key enzymes in the MVA pathway and catalyzes the reduction of HMG-CoA to mevalonate (Friesen & Rodwell, 2004). HMGR is known as main regulator and rate-limiting enzyme in early biosynthesis of sterol and non-sterol isoprenoids in MVA-harboring eukaryotic cells and it is highly regulated at the transcriptional, translational, and post-translational levels (Burg & Espenshade, 2011). In yeast and mammals, HMGR contains a sterol sensing domain (SSD) that is responsible for detecting sterol levels in the cell and maintaining sterol homeostasis (Espenshade & Hughes, 2007). The SSD is located in the N-terminal membrane binding domain of the HMGR enzyme (Burg & Espenshade, 2011). Moreover, it has been reported that genetic manipulations on HMGR, including truncation of N-terminal domain, led to considerable accumulation of terpenes in transgenic plants and yeast (Bansal *et al*., 2018; Bröker *et al*., 2018; Lee *et al*., 2019). Although HMGR has been extensively characterized in model eukaryotic organisms, little is known about its features in diatoms.

**Figure 1:**
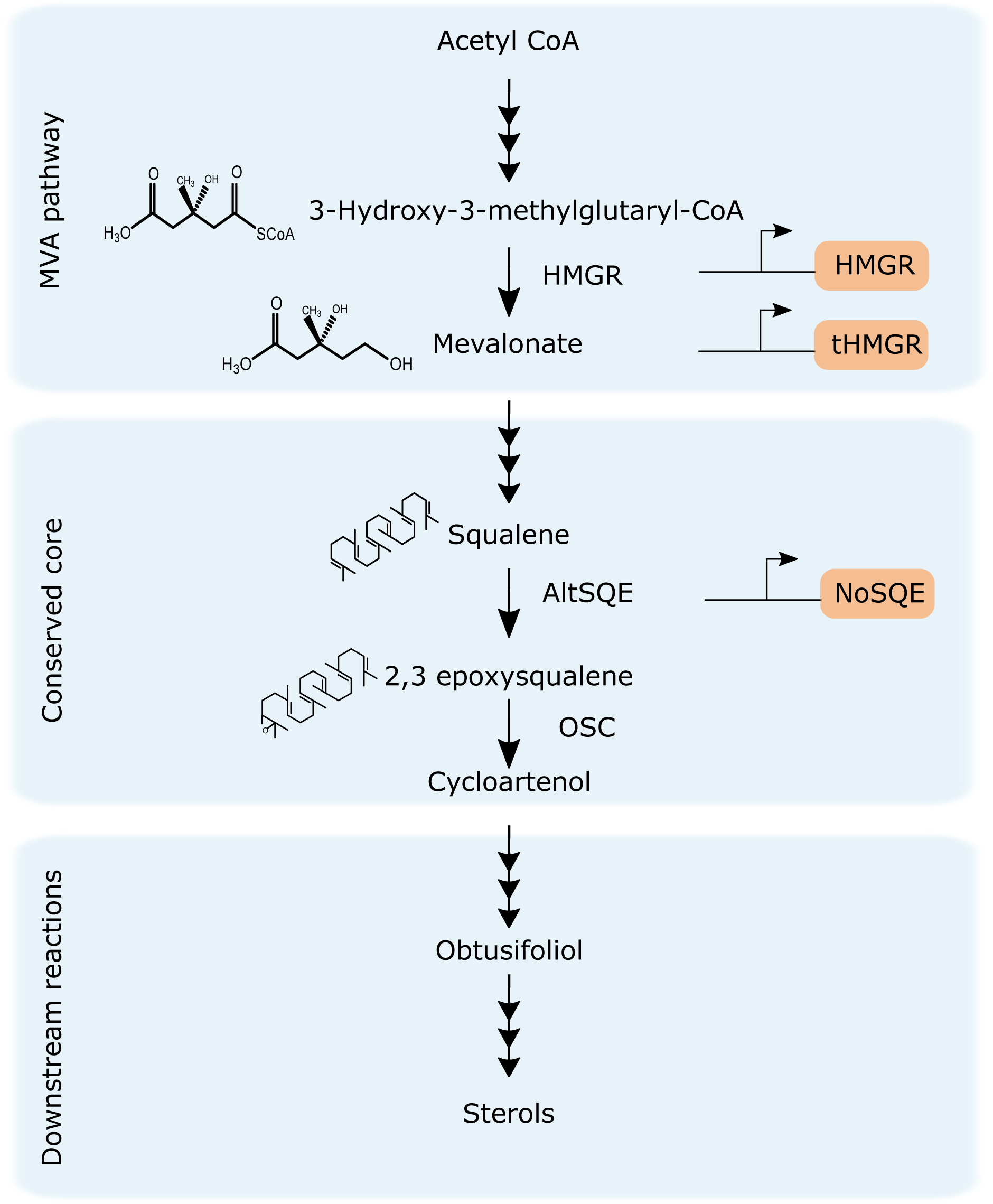
Upstream reactions and conserved core of sterol biosynthesis pathway in diatoms and genetic targets overexpressed in this study, highlighted in orange. Mevalonate pathway, MVA; 3-hydroxy-3-methylglutaryl-coenzyme A reductase, HMGR, truncated HMGR, tHMGR, alternative squalene epoxidase, AltSQE, squalene epoxidase from *N. oceanica*, NoSQE, oxidosqualene cyclase, OSC.

MVA products IPP and DMAPP are subsequently used for the synthesis of squalene, the first committed intermediate in the formation of sterols (Gill *et al*., 2011) (Fig. 1). In plants, fungi, and animals, squalene is converted into 2,3 epoxysqualene. This reaction is conventionally catalyzed by the enzyme squalene epoxidase (SQE, E.C. 1.14.14.17) (Gill *et al*., 2011). Several studies indicate that SQE is a control point in cholesterol synthesis modulated by sterol levels and post-translationally regulated by cholesterol-dependent proteasomal degradation (Nagai *et al*., 2002b; Gill *et al*., 2011). However, diatoms do not possess a conventional SQE, and instead this step is catalyzed by a recently characterized alternative squalene epoxidase (AltSQE) (Pollier *et al*., 2019). The diatom AltSQE belongs to the fatty acid hydroxylase superfamily, and differs from the conventional flavoprotein SQE. Whether AltSQE has a similar role to SQE in sterol regulation is not known.

While selective inhibitors for AltSQE are not known, the treatment of diatoms with statins, known to inhibit *HMGR* enzymes, has resulted in perturbation of isoprenoids metabolism that included an overall decrease of total sterols content (Massé *et al*., 2004; Conte *et al*., 2018), suggesting that a conventional *HMGR* enzyme might be involved in the pathway. In this work, we genetically targeted the HMG CoA reduction and squalene epoxidation steps in *P. tricornutum* to provide further insights into the nature of the diatom sterol biosynthesis pathway and its regulatory constraints. Considering the demonstrated challenges in genetically down-regulating the essential genes involved in *P. tricornutum* sterol metabolism (Pollier *et al*., 2019, Fabris *et al*., 2014), we chose to investigate these pathway nodes by gene overexpression and subcellular localisation.

We generated independent diatom exconjugant lines constitutively expressing (i) either the full-length *HMGR*, (ii) an N-terminal truncated version of *HMGR* (*tHMGR*), or (iii) a heterologous SQE from *Nannochloropsis oceanica* (*NoSQE*) over the background of the endogenous AltSQE. By phenotyping these transgenic diatom cell lines, we describe specific changes in several nodes of the sterol biosynthesis pathway and provide evidence for regulatory mechanisms unique to diatom sterol metabolism.

## METHODOLOGY

### Diatom culturing

The species *P. tricornutum* (CCMP632) and *T. pseudonana* (CCMP1335) were obtained from the National Centre for Marine Algae and Microbiota at Bigelow Laboratory (USA). Axenic cultures were maintained in L1 medium (Guillard & Hargraves, 1993) at 18°C under continuous cool white light (100 µmol photons m^-2^ s^-1^) in a shaking incubator (100 rpm).

### Episome construction and transformation

All episomes used in this study were assembled using uLoop assembly method (Pollak *et al*., 2019). Individual components for episome assembly (L0 parts) were built and domesticated using uLoop assembly syntax. Assembly reactions were performed using the respective uLoop assembly backbones for each level as described by Pollak et al. (2018). After domestication, each L0 part was confirmed by Sanger sequencing. Correct episome assemblies were confirmed by colony PCR and diagnostic restriction digestion. The source of each DNA part and primers used for domestication are listed in Table S1. All L0 parts used to assemble the plasmids used in this work have been deposit in Addgene (Table S1). Plasmid maps (Fig. S1, S2) and complete plasmid sequence are provided in supplemental material.

Episomes consisted of a pCA-derived backbone (Pollak *et al*., 2019), *CEN/ARS/HIS* and *OriT* sequences, a selection cassette, and an expression cassette (Fig. S1, S2). Sequence *OriT* required for bacterial conjugation were amplified from *pPtPBR11* plasmid (Diner *et al*., 2016) (Genebank KX523203). Selection markers nourseothricin (*NAT*) for *T. pseudonana* and blasticidin-S deaminase (BSD) for *P. tricornutum* were driven by elongation factor 2 (*EF2*) constitutive promoters from corresponding diatom species (*T. pseudonana* v. 3 ID: *269148*; *P. tricornutum* v. 3 ID: *Phatr3_J35766*). Expression cassettes included genes encoding either putative *HMGR, tHMGR*, or *NoSQE* each fused at the C-terminus with a mVenus fluorescent protein (Nagai *et al*., 2002a), and an expression cassette expressing only mVenus was used to assemble an empty control vector (Figure S1). The open reading frames encoding putative *HMGR* and *tHMGR* were amplified from genomic DNA of either *T. pseudonana* (CCMP1335) (Gene ID *Thaps_33680*) or *P. tricornutum* (CCMP632) (Gene ID *Phatr3_J16870*) (Table S1). A domesticated, codon-optimized synthetic gene encoding SQE sequence from *Nannochloropsis oceanica* v.2 CCMP1779 (Gene ID *521007*) was obtained from Genewiz® (USA). Expression of target genes were driven by the promoter of elongation factor 2 (*EF2*) in *T. pseudonana* and the promoter of predicted protein *Phatr3_J49202* in *P. tricornutum*. L0 parts for *CEN/ARS/HIS*, fluorescent reporter gene *mVenus, Phatr3_J49202* promoter and terminator were obtained from Dr. Christopher Dupont (J. Craig Venter Institute, USA). The plasmid pTA-Mob for conjugation (Strand *et al*., 2014) was obtained from Dr. Ian Monk (University of Melbourne, Australia).

### Diatom transformation and screening

Diatoms were transformed by bacterial conjugation (Karas et al., 2016). The transformation protocol for *T. pseudonana* was modified by increasing the starting bacterial density (OD_600_ to 0.5) and the final incubation and recovery period for transformed diatom culture to 24 hrs prior to selection on plates containing nourseothricin. *T. pseudonana* and *P. tricornutum* colonies resistant to nourseothricin (50 µg ml-^1^) or blasticidin (10 µg ml^-1^), respectively, were inoculated in 96-multiwell plates containing 200 µl of L1 medium with 100 µg ml^-1^ of nourseothricin or 10 µg ml^-1^ blasticidin, depending on the diatom species, and subcultured every 5 days. Clonal lines from 96-well plates were screened by detecting mVenus fluorescence using a CytoFLEX S (Beckman Coulter) flow cytometer operated in plate mode. 48 clones of each expression system were screened for *T. pseudonana* and 12 for *P. tricornutum*. A 488 nm laser was used for fluorescence excitation; mVenus fluorescence was detected using a 525/40 nm filter and chlorophyll fluorescence was detected using 690/50 nm filter. 10,000 events were analyzed per sample. Three independent cell lines per construct with the highest median mVenus fluorescence readings were selected for full-scale experiments, including WT and empty vector controls.

### Experiments with transgenic diatom cultures

Three replicates of each selected clone were inoculated in 5 ml of L1 medium (100 µg ml^-1^ nourseothricin or 10 µg ml^-1^ blasticidin) and grown for 3 days. Subsequently, cultures were upscaled to 50 ml L1 supplemented with the respective antibiotic for 5 days and these were used to inoculate cultures in L1 medium for sterol analysis experiments. Full-scale experiments were carried out in 200 ml flasks containing 120 ml of L1 medium and antibiotic under continuous light (150 µmol photons m^-2^ s^-1^) and constant shaking (95 rpm). Cell density and mVenus fluorescence were monitored daily by sampling 200 µl from each culture and transferring it to a 96 well plate for high-throughput flow cytometry analysis. Pulse Amplitude Modulated (PAM) fluorometry was used to estimate photosynthetic activity by comparing fluorescence yield of PSII under ambient irradiance (F) and after application of a saturating pulse (Fm) (Schreiber, 2004). After 48 hours of growth, biomass was harvested by centrifuging at 4000 g for 10 minutes. Diatom pellets were washed with Milli-Q water to eliminate excess salt, freeze-dried to determine dry weight, and kept at −20°C until sterol extraction.

### Extraction and analysis of sterols by GC-MS

For sterol extraction, dry cell matter was heated in 1 mL of 10% KOH ethanolic solution at 90°C for one hour. Sterols were extracted from cooled material in three volumes of 400 µL of hexane. An internal standard, 5a-cholestane, was added to each sample. Hexane fractions were dried under a gentle N2 stream, and derivatized with 50 µL of 99% BSTFA + 1% TMCS (N,O-Bis(trimethylsilyl)trifluoroacetamide, Trimethylsilyl chloride) at 70°C for one hour. The resulting extractions were resuspended in 50 µL of fresh hexane prior to GC-MS injection.

Gas chromatography/mass spectrometry (GC-MS) analysis was performed using an Agilent 7890 instrument equipped with a HP-5 capillary column (30 m; 0.25 mm inner diameter, film thickness 0.25 μm) coupled to an Agilent quadrupole MS (5975 N) instrument. The following settings were used: oven temperature initially set to 50°C, with a gradient from 50°C to 250°C (15.0°C min^-1^), and then from 250°C to 310°C (8°C min^-1^, hold 10 min); injector temperature = 250°C; carrier gas helium flow = 0.9 mL min^-1^. A split-less mode of injection was used, with a purge time of 1 min and an injection volume of 5 μL. Mass spectrometer operating conditions were as follows: ion source temperature 230°C; quadrupole temperature 150°C; accelerating voltage 200 eV higher than the manual tune; and ionization voltage 70 eV. Full scanning mode with a range from 50 to 650 Dalton was used.

Sterol peaks were identified based on retention time, mass spectrum, and representative fragment ions compared to the retention times and mass spectrum of authentic standards. The NIST (National Institute of Standards and Technology) library was also used as reference. The area of the peaks and deconvolution analysis was carried out using the default settings of the Automated Mass Spectral Deconvolution and Identification System AMDIS software (v2.6, NIST). Peak area measurements were normalized by both the weight of dry matter prior to extraction, and the within-sample peak area of the internal standard 5a-cholestane. Sterol standards used to calibrate and identify GC-MS results in this study included: cholest-5-en-3-β-ol (cholesterol); (22E)-stigmasta-5,22-dien-3β-ol (stigmasterol); stigmast-5-en-3-β-ol (sitosterol); campest-5-en-3-β-ol (campesterol); (22E)-ergosta-5,22-dien-3-β-ol (brassicasterol); (24E)-stigmasta-5,24-dien-3β-ol (fucosterol); 9,19-Cyclo-24-lanosten-3β-ol (cycloartenol);, 5-α-cholestane; and the derivatization reagent bis(trimethyl-silyl) trifluoroacetamide and trimethylchlorosilane (99% BSTFA + 1% TMCS) and were obtained from Sigma-Aldrich, Australia.

### Fluorescence imaging

Live diatom transformants expressing mVenus were imaged without fixative with a confocal laser scanning microscope (Nikon A1 Plus, Japan) and photomultiplier tube (PMT) detector. The 488-nm and 637-nm lasers were used for mVenus and chlorophyll autofluorescence, respectively. Gains on the detector were kept constant between samples and controls. Images were acquired with 60×/1.4 objective oil immersion objective and processed using imaging software NIS-Elements Viewer 4.0 (Nikon, Japan).

### Multiple sequence alignment and phylogenetic reconstruction

Diatom homologue sequences were retrieved either from the Marine Microbial Eukaryote Transcriptome Sequencing Project (MMETSP) (Keeling *et al*., 2014; Johnson *et al*., 2019) database or from GenBank protein database (Table S2) using BLASTp search with *T. pseudonana HMGR* as the query sequence. The species names and corresponding MMETSP ID numbers are listed in Table S2. *HMGR* from yeast, mammals, and plants were used as outgroups. Sequences from outgroups (Table S2) were obtained from GenBank protein database (complete sequence in Supplementary Material). Multiple sequence alignments of the full-length protein sequences were performed by MAFFT version 7 program with default parameters and alignments were manually edited by exclusion of ambiguously aligned regions. The maximum likelihood tree phylogenetic tree was constructed using MEGA 6 with partial deletion option. The reliability of obtained phylogenetic tree was tested using bootstrapping with 1000 replicates. Prediction of transmembrane helices in *HMGR* from diatoms was carried out using the TMHMM Server v. 2.0 with default parameters (Krogh *et al*., 2001). Conserved motifs in the selected sequences were identified by an InterProScan (Jones *et al*., 2014) search against all available member databases, including Pfam (protein families) and SUPERFAMILY (structural domains).

### Statistical analysis

All plots were generated using R: A language and environment for statistical computing. All experiments were conducted in triplicate. The analyses performed were Shapiro-Wilk to test normality, non-parametric Kruskal–Wallis test and Pairwise Wilcoxon Rank Sum Tests to calculate pairwise comparisons between group levels with corrections for multiple testing. Differences between groups were considered significant at *p* < 0.05.

## RESULTS

### Identification of putative *HMGR* from *T. pseudonana* and *P. tricornutum*

In contrast to the recently discovered AltSQE, little is known about the diatom HMGR enzyme despite being a major regulatory step in sterol biosynthesis. Previous studies demonstrated that specific HMGR inhibitors alter isoprenoids metabolism in the diatoms *P. tricornutum, Haslea ostrearia* and *Rhizosolenia setigera* (Massé *et al*., 2004; Conte *et al*., 2018). Given the presence of many unusual features in diatoms metabolism (Fabris *et al*., 2012; Pollier *et al*., 2019; Jaramillo-Madrid *et al*., 2020a) we analyzed the conservation of the HMGR sequence among all the diatom species with genomic or transcriptomics sequences available.

The amino acid sequence of *Arabidopsis thaliana* HMGR enzyme (AT1G76490) was used as query to search against the genome sequence of the diatoms *T. pseudonana* and *P. tricornutum* to identify the genes putatively encoding *HMGR*. Through this analysis, we identified a single copy of a putative *HMGR* gene located in chromosome 29 in *P. tricornutum* (Gene ID *Phatr3_J16870*)(Fabris *et al*., 2014) and chromosome 4 in *T. pseudonana* (Gene ID Thaps_33680). In model eukaryotic organisms, HMGR is characterized by the presence of an N-terminal membrane domain and a C-terminal catalytic region. Sequence alignment analysis revealed differences in membrane domain location among model organisms, while the catalytic region is conserved (Fig. 2, S3). The C-terminal catalytic domain of HMGR was highly conserved across all the organisms analyzed (Fig. 2). The catalytic residues Glu559, Asp767, and His866 (Friesen & Rodwell, 2004; Li *et al*., 2014), were also found to be present and conserved in *T. pseudonana* and *P. tricornutum* (Fig. S3). HMGRs from *Saccharomyces cerevisiae* and *Homo sapiens* possess a sterol sensing domain (SSD) in the transmembrane N-terminal region, which is involved in sterol homeostasis (Burg & Espenshade, 2011), therefore, we analyzed HMGR transmembrane region in diatoms to identify similarities with other model organisms. Most of the analyzed HMGR sequences from diatoms possess three trans-membrane helices in the N-terminal domain, except a few with two or none domains predicted (Table S2). In comparison, plants usually have two domains (Li *et al*., 2014). We found seven transmembrane domains in *S. cerevisiae* and five in *Homo sapiens* (Table S2). We did not detect similarities with known yeast and mammals SSDs in the N-terminal region in any of the diatoms analyzed (Fig. 2).

**Figure 2:**
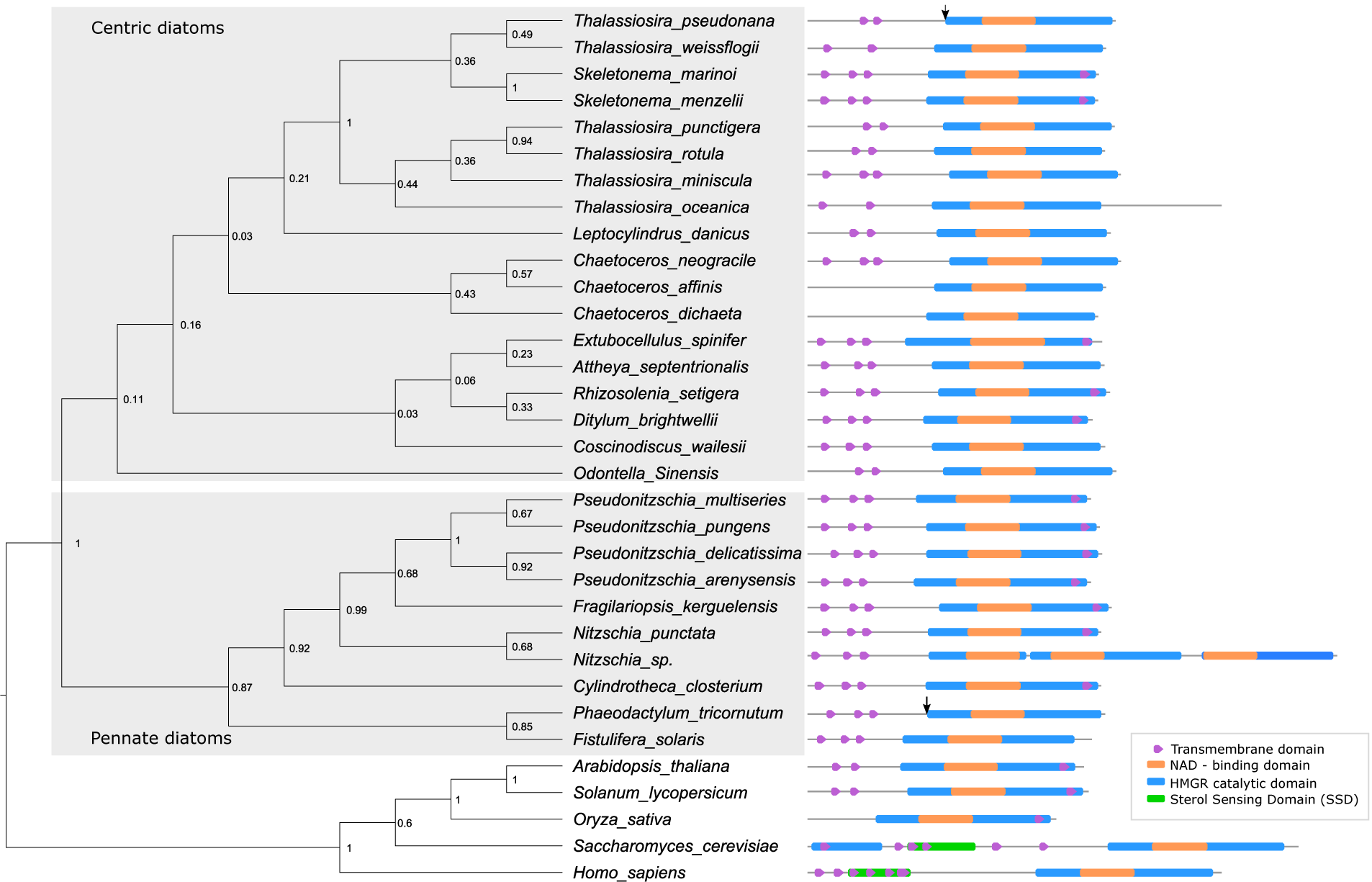
Maximum likelihood phylogenetic tree of diatom HMGR proteins and its domains. Numbers at the nodes represent bootstrap support (1000 replicates). HMGR from yeast, mammals and plants were used as outgroups. Arrow represent start of N-terminal truncated version for *P. tricornutum* and *T. pseudonana* used in this study.

### Phylogenetic analysis of HMGR and conserved protein domains

Based on the alignments of full-length HMGR protein sequences of twenty-eight diatom species retrieved from whole genome and transcriptome assemblies, a maximum likelihood phylogenetic tree was constructed to study evolutionary relationship of HMGR protein sequence among diatoms (Fig. 2). We designated HMG*R* from yeast, mammals, and plants as outgroups. Pennate and centric diatoms were divided into two different clades (Fig. 2). As expected, HMGR from diatoms of the same genus tended to cluster together. Species from the order *Thalassiosirales* which includes *Thalassiosira* and *Skeletonema* genus are grouped together (Fig. 2). Similarly, HMGR from the diatoms *P. tricornutum* and *Fistulifera solaris* that belong to the *Naviculales* order appear to be closely related (Fig. 2). Interestingly, we did not find a match for HMGR in the transcriptomic sequences of the diatoms: *Chaetoceros muelleri*, as previously reported (Jaramillo-Madrid *et al*., 2020a), *Chaetoceros brevis, Chaetoceros debilis* and *Chaetoceros curvisetus* (Table S2).

### Expression and subcellular localization of putative HMGR and tHMGR

While the core reactions in sterol synthesis being conserved in *T. pseudonana* and *P. tricornutum* (Jaramillo-Madrid *et al*., 2020a), both diatoms produce a distinctive profile of sterol compounds which variates differently upon changing environmental conditions and chemical inhibitors treatment (Jaramillo-Madrid *et al*., 2020a,b). Additionally, the sterol metabolism of the centric diatom *T. pseudonana* has not been explored to the same depth as the model pennate *P. tricornutum*. To investigate the subcellular localization and evaluate the effect of overexpression of the rate-limiting enzyme HMGR on sterol accumulation, *T. pseudonana* and *P. tricornutum* were transformed with episomes containing their respective putative *HMGR* copy fused to mVenus, driven by a constitutive promoter (Fig. 1, Fig. S1,S2). Episomes are maintained extra-chromosomally and therefore enable more consistent expression required for metabolic studies (George *et al*., 2020). The trans-membrane domains of HMGR enzymes in mammals and yeast have been reported to contain a sterol sensing domain (SSD) that regulates expression and degradation of the enzyme (Kuwabara & Labouesse, 2002). Although our results suggest that diatom HMGRs lacks a SSD (Fig. 2), we designed an N-terminal truncated version, tHMGR, to evaluate whether an unknown regulatory sequence is present in N-terminal region affecting activity of HMGR in diatoms. This *tHMGR* sequence encoded solely the C-terminal catalytically active region of the enzyme.

The mVenus fluorescence was measured by flow cytometry and used as an indirect proxy of enzyme expression, since each expression system was C-terminal fused with mVenus protein. Three clones per expression system with the highest median mVenus signal were chosen for full scale experiments. In *P. tricornutum*, time course median mVenus fluorescence in mVenus control clones was 10-fold compared to WT, which confirms the effectiveness of the chosen promoter (Fig. S4). Conversely, median mVenus fluorescence clones expressing HMGR-mVenus and tHMGR-mVenus was 1.3-fold compared to WT, indicating an apparent regulation process occurring over the fused proteins (Fig. S4). In *T. pseudonana*, median mVenus fluorescence in HMGR-mVenus, tHMGR-mVenus, and mVenus control clones appear similar to WT signal, suggesting that low expression was achieved using the *EF2* promoter (Fig. S5). However, confocal microscopy images confirmed expression of mVenus in both diatom species (Fig. 3). Different cellular localizations were observed for each genetic construct. Images of control cell lines showed mVenus expression localized in the cytoplasm (Fig. 3), while no mVenus fluorescence was detected in WT diatoms. mVenus fluorescence in exconjugants overexpressing HMGR-mVenus was detected around the chloroplast, suggesting that putative HMGR is localized in the endoplasmic reticulum (ER), which tightly surrounds the chloroplasts in diatoms (Kroth, 2002) (Liu *et al*., 2016). Conversely, tHMGR-mVenus localises in the cytoplasm, consistent with truncation of the N-terminal membrane domain (Fig. 3).

**Figure 3:**
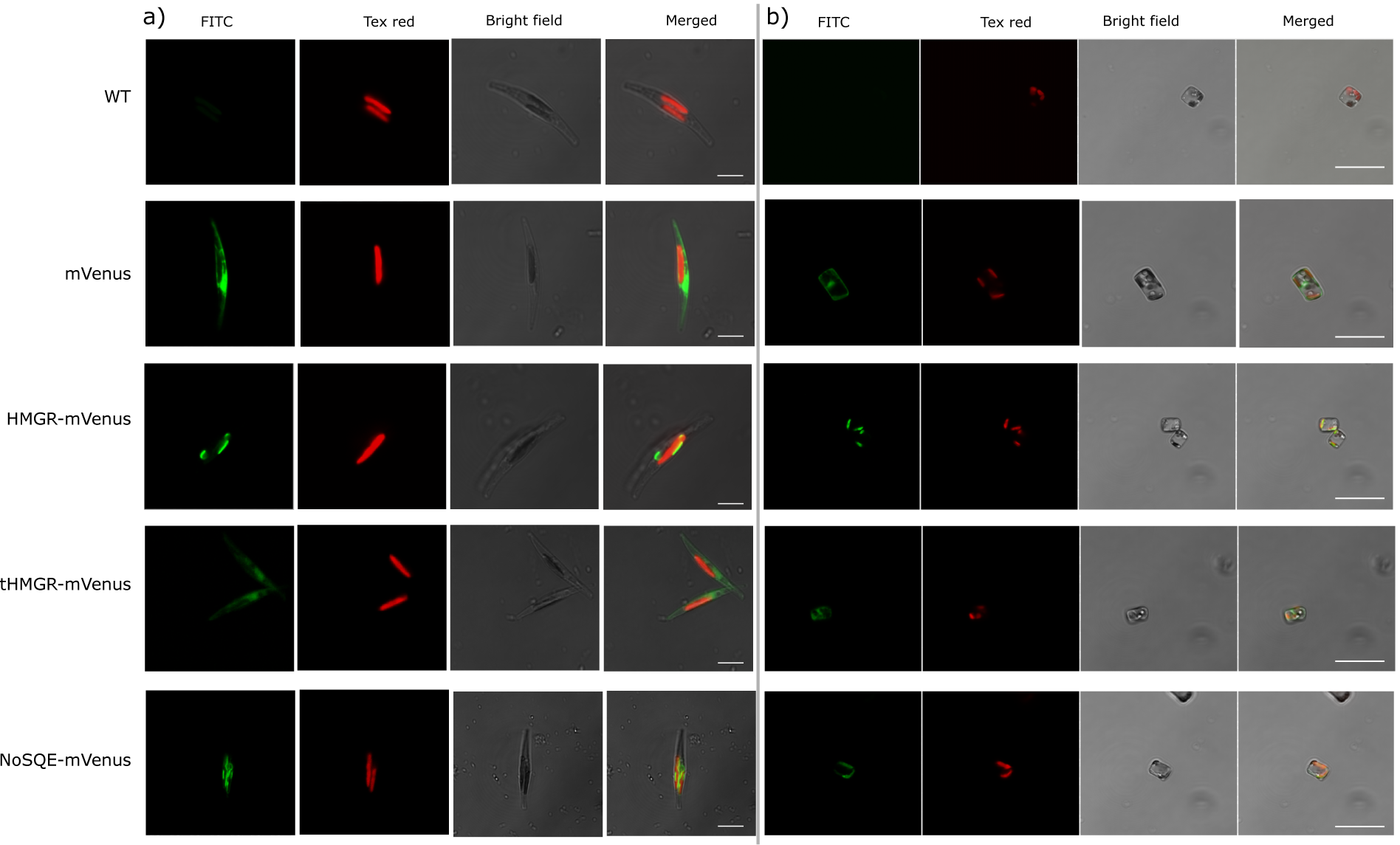
Confocal microscopy images showing subcellular localization of the mVenus fusion with target proteins in representative transgenic a) *P. tricornutum* cells, scale bars correspond to 5 *µ*m and b) *T. pseudonana* cells, scale bars correspond to 10 *µ*m. Wild type (WT) as negative control and control cell lines that only expressed mVenus. 3-hydroxy-3-methylglutaryl-coenzyme A reductase, HMGR-mVenus, truncated HMGR, tHMGR-mVenus, squalene epoxidase from *N. oceanica*, NoSQE-mVenus. Scale bars correspond to 10 *µ*m.

### Influence of HMGR and tHMGR expression on sterol levels in *T. pseudonana* and *P. tricornutum*

Although expression of transgenes appears to be low according to mVenus fluorescence levels, we proceeded to identify changes in sterol profiles in the transgenic lines. Sterols were extracted from exconjugants in the mid-exponential phase, which was the time period with the maximum observed mVenus fluorescence (Fig. S4,S5) with enough biomass to sample for sterol extraction (determined to be 48 hours growth for *T. pseudonana* and 72 hours for *P. tricornutum*). After 75 hours, cell density of *P. tricornutum* HMGR-mVenus and tHMG-mVenus R was 1.4 times lower than WT, while no growth impairment was observed for mVenus exconjugants (Fig. S6). No differences in chlorophyll levels and in effective quantum yield of PSII were observed in *P. tricornutum* exconjugants (Fig. S7, S8). Similarly, no growth impairment, chlorophyll levels or differences in effective quantum yield of PSII compared to WT were observed for *T. pseudonana* exconjugants (Fig. S9, S10, S11).

In *P. tricornutum* overexpressing HMGR-mVenus, squalene levels were ten times higher in HMGR-mVenus exconjugants than in WT and mVenus controls. Moreover, a 3-fold increase in cycloartenol and a 2.5-fold obtusifoliol accumulation was detected compared to WT (Fig. 4). However, we did not detect the intermediate 2,3 epoxysqualene. Levels of end-point sterol campesterol decreased 2-fold, whereas no significant differences were observed in brassicasterol, the most abundant sterol in *P. tricornutum*. We detected traces levels of end point sterol 24-methylcholesta-5,24(24’)-dien-3β-ol in the WT and mVenus controls, which is typically found in centric diatoms and has not been reported in *P. tricornutum* (Rampen *et al*., 2010). Levels of this end-point sterol were 17 times higher in two independent exconjugant lines expressing HMGR-mVenus. Total sterol levels were not affected despite significant changes in individual sterols (Fig. 4).

**Figure 4:**
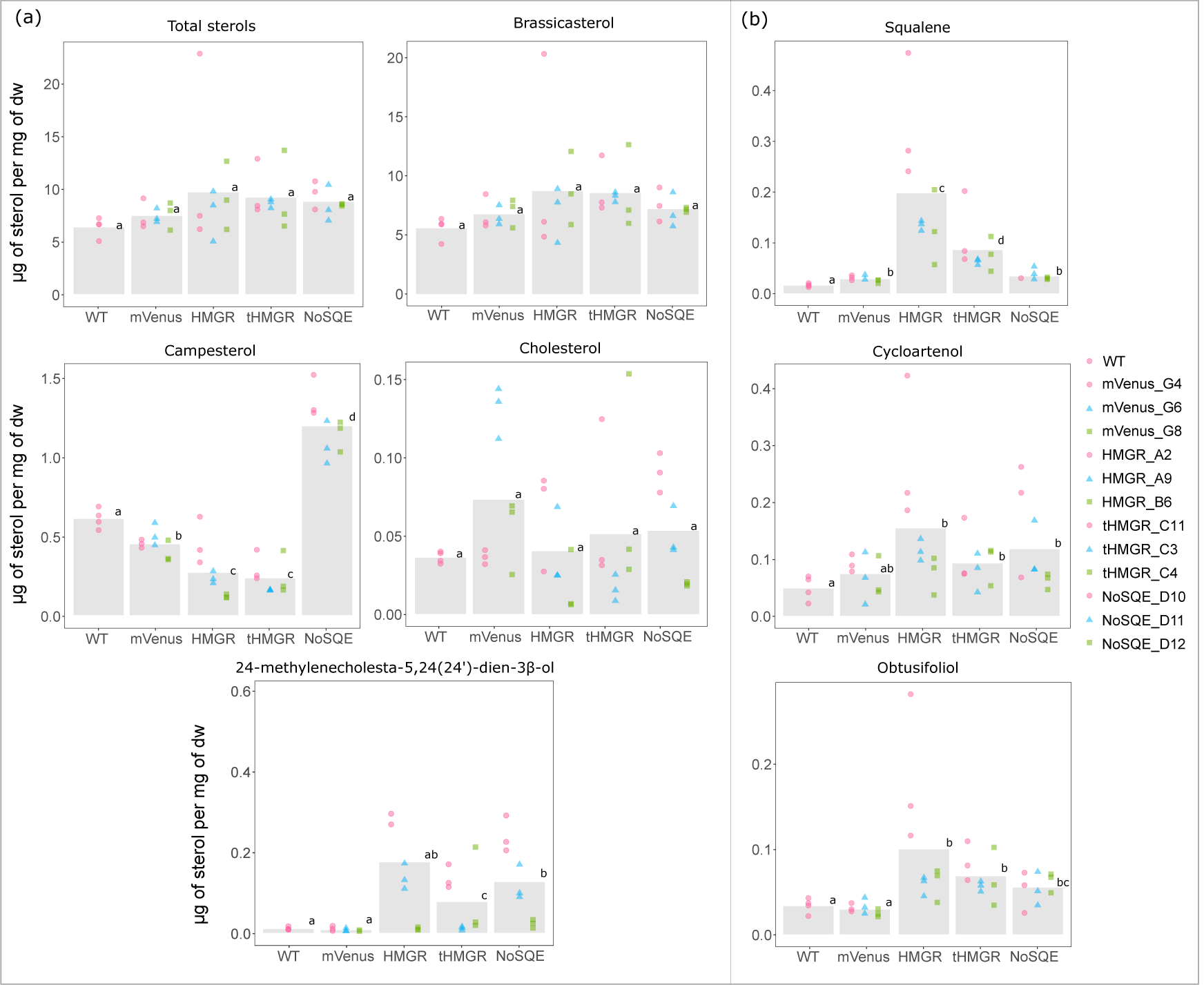
Sterol levels in *P. tricornutum* transformants. (a) End-point sterol (b) intermediates accumulation. Identical letters denote no statistically significant differences among groups using the Pairwise Wilcoxon Rank Sum tests (p < 0.05, n = 9).

Since expression of mVenus, HMGR-mVenus, and tHMGR-mVenus in *T. pseudonana* did not appear to be effective based on flow cytometry data (Fig. S5), observed changes on sterol profiles may not be directly related to the overexpression of the targeted enzymes. *T. pseudonana* cell lines transformed with *HMGR*-mVenus construct exhibited a decrease in the minor sterols fucosterol and isofucosterol relative to WT control (Fig. S12). However, isofucosterol reduction was also detected in the control expressing only mVenus (Fig. S12). No intermediates were detected. Total sterol levels remained similar in the three independent cell lines studied; no significant differences were observed in comparison to the WT control (Fig. S12).

Expressing the catalytically active region of their putative HMGRs (tHMGR) was expected to reduce regulatory mechanisms that may affect HMGR activity in diatoms. In *P. tricornutum*, sterol changes in cell lines expressing tHMGR-mVenus were similar to those expressing HMGR-mVenus, with campesterol levels 2-fold less abundant than in HMGR-mVenus transformants (Fig. 4). Traces of 24-methylcholesta-5,24(24’)-dien-3β-ol were detected as in HMGR-mVenus expressing lines, 10 times higher compared to WT. We also detected an increase in the intermediates squalene (4-fold), cycloartenol (1.8-fold) and obtusifoliol (2-fold) compared to WT (Fig. 4). Squalene levels accumulated in HMGR-mVenus clones were statistically different to tHMGR clones, being 2.5 times higher in HMGR-mVenus expressing lines (Fig. 4). No changes in total sterol levels and brassicasterol were observed (Fig. 4).

No changes in total sterol content were observed in *T. pseudonana* cell lines transformed with *tHMGR*-mVenus (Fig. S12). No intermediates were detected. Significant changes in less abundant sterol compounds occurred in transformants including controls expressing only mVenus.

### Heterologous expression of a Stramenopile putative SQE

Diatoms have been reported to employ an alternative squalene epoxidase (AltSQE) that is different from the conventional SQE found in other eukaryotes (Pollier *et al*., 2019). It has been observed that artificially alter the expression of this enzyme in *P. tricornutum* is particularly challenging (Pollier *et al*., 2019), suggesting that a strict regulation of endogenous AltSQE may be occurring as is the case for SQE in (Nagai *et al*., 2002b; Gill *et al*., 2011). Therefore, we hypothesized that the expression of a heterologous, conventional SQE that could override endogenous regulation, would influence final sterol levels. Consequently, a heterologous putative SQE from the Stramenopile *N. oceanica* (*NoSQE*, Nanoce ID 521007) was expressed in the diatoms *T. pseudonana* and *P. tricornutum*.

Confocal microscopy images showed that mVenus fluorescence in NoSQE–mVenus transformants was similarly located to HMGR-mVenus, indicating ER localization in both diatom species (Fig. 3). Chlorophyll and mVenus fluorescence intensity were comparable to those of diatoms overexpressing putative HMRG-mVenus and tHMGR-mVenus (see 3.2 section) (Fig. S4, S5, S7, S10).

Total sterol content of *T. pseudonana* and *P. tricornutum* exconjugants transformed with *NoSQE*-mVenus constructs remained unchanged compared to WT and mVenus controls (Fig. 4, S12). However, in *P. tricornutum* overexpressing NoSQE-mVenus we observed significant differences in both end-point sterols and sterol intermediates. Downstream intermediates cycloartenol and obtusifoliol exhibited a 1.8-fold increase compared to WT control (Fig. 4), while no differences in squalene were observed. The intermediate 2,3 epoxysqualene was not detected in either diatom species. In contrast to HMGR-mVenus overexpression, campesterol increased by 3-fold (Fig. 4). However, the major end-point sterol brassicasterol remained unchanged.

## DISCUSSION

### HMGR is largely conserved among diatoms and lacks a conventional sterol sensing domain

To investigate sequence characteristics of the rate-limiting HMGR enzyme in diatoms, we identified the genes putatively encoding the enzyme HMGR from 28 different diatom species. While a putative HMGR homologue was detected in 28 of the diatom species, the failure to detect HMGR transcripts in transcriptomic sequences of some diatoms belonging to *Chaetoceros* genus (Table S2), may support the hypothesis that that those diatoms may solely rely on the MEP pathway to produce isoprenoids (Jaramillo-Madrid *et al*., 2020a). The lack of obvious HMGR transcripts may also have occurred due to low expression or down-regulation of these and other genes related to the MVA pathway under the conditions in which RNA sequencing was performed. Nevertheless, conserved HMGR genes were detected in many diatoms for which genomic or transcriptomic data are available. These HMGR genes diverge between pennate and centric diatoms (Fig. 2), which are separated by 90 million years of divergent evolution (Bowler *et al*., 2008). Presence of putative HMGR in most of the diatom species analysed is an indicator of presence of MVA pathway operating in diatoms, as it was previously reported by transcriptomics analysis in the diatoms *P. tricornutum* and *T. pseudonana* (Jaramillo-Madrid *et al*., 2020a). The MEP pathway appears functional in the two model diatoms, which indicates that both cytosolic MVA and plastidial MEP are simultaneously operating in *P. tricornutum* and *T. pseudonana*, as it is the case in plants (Vranova *et al*., 2013; Jaramillo-Madrid *et al*., 2020a). Presence of both pathways could represent an advantage for terpenoids production, due to a potentially higher metabolic flux and greater pool of available precursors (Vavitsas *et al*., 2018).

The organization of the N-terminal transmembrane domain of diatom HMGR differs significantly to their mammals and yeast counterparts. While most of the diatoms analyzed in this study presented three transmembrane domains (Fig. 2, Table S2), mammals possess five and yeast seven. The presence of transmembrane domains is likely related to anchoring the protein within the ER membrane, but the consequences of this structural difference for diatom HMGR in terms of regulation of enzyme expression and activity are unknown. In mammals and yeast, HMGR possess a SSD involved in sensing oxysterol molecules that activate feedback regulation leading to degradation of the protein (Burg & Espenshade, 2011) (Theesfeld *et al*., 2011). Despite the lack of a conventional sterol-sensing domain in the HMGR enzymes of diatoms (Fig. 2), several studies have shown a transcriptional response of MVA enzymes to perturbations in sterol metabolism (D’Adamo *et al*., 2019)(Jaramillo-Madrid *et al*., 2020a). The fact that in diatom sequences do not possess a canonical SSD opens several possibilities, including that HMGR may not play the same regulatory role in diatoms as it does in other organisms; i.e. the MVA pathway may be regulated through a different mechanism that does not involve *HMGR* feedback regulation. Another possible explanation is that HMGRs of diatoms and plants possess a non-conventional SSD sequence, a motif with different characteristics than the already described in mammals and yeast.

### HMGR overexpression lead to accumulation of sterol pathway intermediates in *P. tricornutum*

In this study, we investigated the response of diatoms to genetic targeting of sterol biosynthesis, through manipulation of the MVA rate-limiting enzyme HMGR. The fluorescent localization of extra-chromosomally expressed HMGR-mVenus to the membrane surrounding the plastid is consistent with ER localization of proteins, including AltSQE, from previous studies in *P. tricornutum* (Kroth, 2002; Pollier *et al*., 2019) and *T. pseudonana* (Sheppard *et al*., 2010). Therefore, our study provides evidence that diatom HMGR is localized to the ER, just as in mammals, yeast, and higher plants (Leivar *et al*., 2005; Burg & Espenshade, 2011). The truncation of the native HMGR sequence resulted in cytoplasmic localization, demonstrating that the signals for protein targeting are in the N-terminal portion of the protein sequence.

In some cases, overexpression of HMGR has been showed to slightly increase sterol and intermediates production in other organisms. The overexpression of HMGR from the plant *Panax ginseng* in *Arabidopsis thaliana* resulted in a nearly two-fold increase of sitosterol, campesterol, and cycloartenol, while levels of squalene and stigmasterol did not significantly change (Kim *et al*., 2014). In a more recent study, Lange et al. (2015) independently over-expressed all genes participating in the MVA pathway, obtaining a significant increase in total sterols when expressing HMGR (3.4-fold) and 3-hydroxy-3-methylglutaryl-co-enzyme-A synthase, HMGS (2-fold). The overexpression of native HMGR in *Arabidopsis* (*HMG1*) led to high levels of HMGR mRNA, but only a slight increase in HMGR activity, and no changes in leaf sterol levels (Re *et al*., 1995). Heterologous expression of HMGR from *Hevea brasiliensis* in tobacco, however, showed an increase in HMGR transcript and total sterol from leaves (Schaller *et al*., 1995).

In this study, overexpression of endogenous putative HMGR-mVenus in *P. tricornutum* resulted in an increased accumulation of the intermediates squalene, cycloartenol, and obtusifoliol and decrease in the end-point sterol campesterol. However, we did not detect 2,3 epoxysqualene, indicating possible differences on the catalytic rates of the enzymes squalene epoxidase and cycloartenol cyclase (Fig. 1). While accumulation of squalene is detectable, 2,3 epoxysqualene seems to be rapidly converted into cycloartenol. Accumulation of 2,3 epoxysqualene has been reported before by chemically inhibiting cycloartenol cyclase with Ro 48-8071 (Fabris *et al*., 2014; Jaramillo-Madrid *et al*., 2020a).

Accumulation of intermediates suggest that in *P. tricornutum*, overexpression of HMGR-mVenus boosted production of presumably MVA-derived intermediates, IPP and DMAPP that are subsequently converted into squalene (Fig. 1). Even though MVA products also serve as building blocks of other isoprenoids (Lange *et al*., 2000), perturbation of the rate-limiting step catalyzed by HMGR was sufficient to cause accumulation of downstream intermediates committed to sterol biosynthesis, such squalene, cycloartenol, and obtusifoliol. However, this metabolic bottleneck did not translate into overall increase of sterol compounds; on the contrary, levels of the end-point sterol campesterol were reduced and brassicasterol levels remained unchanged.

Although, fluorescence levels in *P. tricornutum* exconjugants expressing the target enzymes were considerably lower compared to the expression of mVenus alone (Fig. S5), phenotypic changes in terms of sterol profiles indicates that the level of expression was sufficient to cause perturbations in the metabolic pathway (Fig. 4). Moreover, since all the enzymes expressed are membrane proteins, as confirmed by confocal microscopy (Fig. 3, S3), fluorescent signal could have been hindered and no direct correlation with expression could be assumed. The absence of detectable sterol pathway intermediates in *T. pseudonana* transformants (Fig. S12) may be related with the promoter chosen for overexpression of the target enzymes, as higher expression may be necessary to trigger the accumulation of intermediates observed in *P. tricornutum*. Fluorescence signal for mVenus control in *P. tricornutum* (*Phatr3_J49202* promoter) was around ten times higher than in *T. pseudonana* mVenus control (*EF2* promoter) throughout the full-scale experiment (Fig. S4, S5), suggesting that use of stronger promoters for metabolic engineering of *T. pseudonana* should be considered. Levels of fucosterol and isofucosterol decreased in *T. pseudonana* lines transformed with a putative endogenous HMGR-mVenus (Fig. S12). However, since similar results were observed for isofucosterol in the control expressing only mVenus, it is uncertain if the observed reduction was a direct consequence of putative HMGR-mVenus overexpression alone. No intermediates were detected in *T. pseudonana* overexpressing putative HMGR-mVenus. End-point sterols 24-methylenecholesta-5,24(24’)-dien-3β-ol, cholesterol, campesterol, and total sterol levels were statistically indistinguishable from those obtained with untransformed WT and mVenus control transformants (Fig. S12).

Results suggest that there are several regulation points in sterol biosynthesis in diatoms, including the MVA pathway, conserved core, and specialized downstream reactions. It is also possible that MEP responds to an alteration on the MVA pathway, rebalancing IPP and DMAPP pools, and metabolic cross-talk between this two pathways could potentially occur in diatoms. Despite the core reactions in sterol synthesis being conserved in *T. pseudonana* and *P. tricornutum* (Jaramillo-Madrid *et al*., 2020a), we observed different responses in lines overexpressing putative *HMGR*-mVenus of these two model diatoms (Fig. 4, S12). As previously mentioned, these results might be related with the promoters chosen for each species, or could indicate differences on sterol regulation between centric and pennate diatoms that correlates with divergence between putative *HMGR* from both diatom groups (Fig. 2). Differences in sterol profiles from *T. pseudonana* and *P. tricornutum* have been suggested to occur in downstream reactions of sterol synthesis (Jaramillo-Madrid *et al*., 2020a), and responses to alteration on a key point of MVA pathway suggest possible divergences in regulation mechanisms.

### Overexpression of tHMGR does not circumvent native regulatory mechanisms

Additional strategies have been developed to overcome regulation by HMGR and increase MVA carbon flux in other eukaryotes. Truncation of HMGR to remove N-terminal membrane and SSD domain was first reported in plants and yeast with the aim to express only the catalytic domain of HMGR and avoid regulatory effects (Donald *et al*., 1997; Polakowski *et al*., 1998). Although HMGRs of plants do not contain an SSD sequence (Fig. 2), expression of an N-terminal truncated HMGR has been reported to increase sterol levels. Expression of tHMGR from hamster in tobacco resulted in augmented sterol content in leaf tissue (Chappell *et al*., 1995). Constitutive expression of tHMGR from *Hevea brasiliensis* in tobacco resulted in an increasing 11-fold of seed HMGR activities and 2.4-fold increase in total seed sterol content (Harker *et al*., 2003). However, overexpression of a tHMGR has not always been effective at altering sterol content; the overexpression of tHMGR in yeast resulted in accumulation of squalene, no changes in ergosterol, the final sterol in yeast (Donald *et al*., 1997; Polakowski *et al*., 1998). Likewise, in this study we did not observe a statistically significant alteration in total sterols of *T. pseudonana* and *P. tricornutum* after constitutive expression of a tHMGR-mVenus (Fig. 4, S12). Yet, accumulation of the intermediates squalene, cycloartenol, and obtusifoliol was observed in *P. tricornutum*. Interestingly, levels of those intermediates were higher when expressing HMGR-mVenus, suggesting that truncation may have affected enzyme activity, performance, or access to substrates. Although we observed changes in intermediates and minor sterol levels in *P. tricornutum* expressing HMGR-mVenus and tHMGR-mVenus, total sterol levels remained unchanged. These results suggest that diatoms have a tight sterol regulation system that may not be related to the conventional regulation model that involves SSD but rather a complex system with several regulation points not only in the MVA pathway but further down the sterol metabolic pathway.

### Levels of end-point campesterol increased after heterologous expression of NoSQE in *P. tricornutum*

Diatoms possess a distinct AltSQE (Pollier *et al*., 2019) catalyzing the conversion of squalene into 2,3 epoxysqualene which is then transformed into cycloartenol, the first committed step towards the production of steroids (Fig. 1). Whether the presence of an AltSQE confers diatoms and other microeukaryotes with specific biological advantages is not yet known. Similarly, AltSQE and SQE are mutually exclusive and, to date, no organisms have been found to naturally harbor both (Pollier *et al*., 2019).

The genetic manipulation of SQE and squalene synthase, SQS, has been extensively used for enhanced production of squalene and triterpenoids (Lee *et al*., 2004; Dong *et al*., 2018) (Gohil *et al*., 2019). Point mutations in the SQE gene (ERG1) in yeast resulted in accumulation of squalene (Garaiova *et al*., 2014). Similarly, accumulation of squalene was observed in the green microalgae *Chlamydomonas reinhardtii* after knocking-down the SQE gene, while co-transformation lines with SQE-overexpression and SQE-knockdown yielded similar amounts of squalene (Kajikawa *et al*., 2015).

This is the first study to investigate the response of diatoms to the expression of a conventional SQE. We did not observe any significant changes in growth and photosynthetic phenotypes of *T. pseudonana* and *P. tricornutum* expressing heterologous NoSQE (Fig. S6-S11). This suggest that in diatoms there is no apparent toxicity or physiological reason for the mutual exclusivity between AltSQE and conventional SQE. Confocal microscopy images of lines expressing NoSQE-mVenus fusion proteins, revealed that heterologous NoSQE was proximal to the chloroplasts, indicating that diatoms could recognize the ER signal peptide on the heterologous NoSQE, localizing it in the ER membrane (Fig. 3, S3), just as the endogenous AltSQE (Pollier et al., 2019) and native SQE enzymes are in other species (Leber *et al*., 1998; Laranjeira *et al*., 2015).

Significant accumulation of cycloartenol (1.8-fold) and obtusifoliol (1.8-fold) intermediates, but not of 2,3 epoxysqualene was obtained for *P. tricornutum* lines expressing NoSQE-mVenus (Fig. 4b). These intermediates occur after the formation of 2,3 epoxysqualene, which is the product of the reaction catalyzed by SQE (Fig. 1). As expected, we did not observe increased accumulation of squalene, which is the substrate for SQE, contrary to accumulation obtained by expressing HMGR-mVenus which is upstream of squalene production (Fig. 1). Nevertheless, heterologous expression of NoSQE-mVenus resulted in a 2-fold increase of campesterol, an end-point sterol. Accumulation of intermediates was higher in *P. tricornutum* cell lines expressing HMGR compared to those expressing NoSQE-mVenus (Fig. 4a). This indicates that intermediates accumulated by MVA pathway manipulation (i.e. HMGR) do not necessarily increase the flux to brassicasterol, suggesting that sterol regulation is occurring at the conserved core point and at other points further down the metabolic pathway. In particular, this might suggest that in *P. tricornutum* the epoxidation of squalene might be involved in pathway flux modulation, as observed in mammals (Nagai *et al*., 2002b; Gill *et al*., 2011), and that, to complete the scenario suggested by the results obtained by expressing NoSQE, it is plausible that an additional pathway checkpoint exists at the level of the C22-desaturation (E.C 1.14.19.41, Phatr3_J51757) (Fabris *et al*., 2014). When treated with fluconazole and fenpropimorph, inhibitors targeting upstream of campesterol, the transcription of *Phatr3_J51757* significantly increases (Jaramillo-Madrid *et al*., 2020a). This further supports the hypothesis that the last reaction in sterol synthesis could be a highly regulated point to maintain stable sterol levels in the cell. These results suggest that to observe changes in end-point sterol compounds, increasing the precursors pool is not enough; genetic manipulation should target other points further down in the metabolic pathway, such as committed steps in sterol synthesis. A future co-expression approach to increase end-point sterol compounds in *P. tricornutum* could involve simultaneous expression of enzymes in the conserved core (i.e. SQE, AltSQE, cycloartenol synthase) and enzymes further down such as sterol C-22 desaturase. Similar co-expression approaches for manipulation of sterol levels in diatoms has not been reported but has been used to increase triterpenoid production in other organisms (Lange *et al*., 2015; Zhang *et al*., 2017; Bröker *et al*., 2018; Dong *et al*., 2018).

## Conclusions

The results obtained in this study demonstrate the effectiveness of extra-chromosomal expression of key enzymes involved in sterol synthesis to influence levels of specific sterol compounds. We confirmed reported advantages of the use of extra-chromosomal episomes transformed via conjugation such as expression consistency among clones (Fig. S4-S11) and no random genome integration (George *et al*., 2020). Additionally, we demonstrated the convenience of a modular assembling systems as uLoop (Pollak *et al*., 2019) to build versatile genetic constructs for a functional genetics study with multiple species.

Furthermore, we applied reproducible genetic transformation methods for extra-chromosomal and heterologous expression to provide important insights into the metabolic bottleneck and pathway-level regulation of sterol synthesis in diatoms. We obtained accumulation of sterol pathway intermediates by overexpression of HMGR-mVenus, indicating possible metabolic bottleneck(s) downstream of the MVA pathway that may limit flux into end-point sterols.

Accumulation of pathway intermediates is interesting from a biotechnological perspective, as an increased intermediate pool could be used by heterologous pathways plugged into endogenous (tri)terpenoid synthesis, allowing production of other high-value terpenoids as geraniol (Fabris *et al*., 2020).

Whilst significant accumulation of intermediates participating in sterol synthesis was observed in *P. tricornutum* transformants, *T. pseudonana* and *P. tricornutum* transformants did not appear to produce different levels of total sterols. It is presumed that several levels of regulation could be affecting the expression, localization, lifetime, and activities of introduced genes. Further research into the regulatory responses of diatoms to heterologous overexpression may provide further insights into these processes and improve strategies for more informed metabolic engineering approaches.

## Acknowledgements

A.C.J.M. was supported by the UTS International Research Scholarship and President’s Scholarship, as well as a Scholarship from the non-profit Colfuturo Foundation of Colombia. J.A. is the recipient of an Australian Research Council Discovery Early Career Award (DE160100615) funded by the Australian Government. M.F. is supported by a CSIRO Synthetic Biology Future Science Fellowship, co-funded by CSIRO and the University of Technology Sydney. We wish to thank Dr. Unnikrishnan Kuzhiumparambil, Taya Lapshina and UTS Microbial Imaging Facility for methodological guidance.

## Author Contributions

A.C.J.M. designed and performed all experiments and wrote the manuscript. J.A. designed, advised and materially supported experiments, performed bioinformatics, and assisted in writing. R.A. advised experiments, reviewed and assisted in writing. M.F. advised experiments, reviewed and assisted in writing. P.J.R. advised and materially supported experiments.

## REFERENCES

Aldini R, Micucci M, Cevenini M, Fato R, Bergamini C, Nanni C, Cont M, Camborata C, Spinozzi S, Montagnani M, et al. 2014. Antiinflammatory effect of phytosterols in experimental murine colitis model: prevention, induction, remission study. PLoS One 9: e108112.

Armbrust EV. 2009. The life of diatoms in the world’s oceans. Nature 459: 185–192.

Athanasakoglou A, Grypioti E, Michailidou S, Ignea C, Makris AM, Kalantidis K, Mass G, Argiriou A, Verret F. 2019. Isoprenoid biosynthesis in the diatom Haslea ostrearia. New Phytologist 222: 230–243.

Bansal S, Narnoliya LK, Mishra B, Chandra M. 2018. HMG-CoA reductase from Camphor Tulsi (Ocimum kilimandscharicum) regulated MVA dependent biosynthesis of diverse terpenoids in homologous and heterologous plant systems. Scientific Reports 8: 1–15.

Borowitzka MA. 2013. High-value products from microalgae — their development and commercialisation. Journal of Applied Phycology 25: 743–756.

Bröker JN, Müller B, van Deenen N, Prüfer D, Gronover CS. 2018. Upregulating the mevalonate pathway and repressing sterol synthesis in Saccharomyces cerevisiae enhances the production of triterpenes. Applied microbiology and biotechnology 102: 6923–6934.

Burg JS, Espenshade PJ. 2011. Regulation of HMG-CoA reductase in mammals and yeast. Progress in Lipid Research 50: 403–410.

Chappell J, Wolf F, Proulx J, Cuellar R, Saunders C. 1995. Is the reaction catalyzed by 3- hydroxy-3-methylglutaryl coenzyme A reductase a rate-limiting step for isoprenoid biosynthesis in plants? Plant Physiology 109: 1337–1343.

Conte M, Lupette J, Seddiki K, Meï C, Dolch L-J, Gros V, Barette C, Rébeillé F, Jouhet J, Maréchal E. 2018. Screening for biologically annotated drugs that trigger triacylglycerol accumulation in the diatom Phaeodactylum. Plant physiology 177: 532–552.

D’Adamo S, Schiano di Visconte G, Lowe G, Szaub-Newton J, Beacham T, Landels A, Allen MJ, Spicer A, Matthijs M. 2019. Engineering the unicellular alga Phaeodactylum tricornutum for high-value plant triterpenoid production. Plant Biotechnology Journal 17: 75–87.

Diner RE, Bielinski VA, Dupont CL, Allen AE, Weyman PD. 2016. Refinement of the diatom episome Maintenance sequence and Improvement of Conjugation-Based dNA delivery Methods. Frontiers in Bioengineering and Biotechnology 4: 1–12.

Donald KA, Hampton RY, Fritz IB. 1997. Effects of overproduction of the catalytic domain of 3-hydroxy-3- methylglutaryl coenzyme A reductase on squalene synthesis in Saccharomyces cerevisiae. Applied and Environmental Microbiology 63: 3341–3344.

Dong L, Pollier J, Bassard J, Ntallas G, Almeida A, Lazaridi E, Khakimov B, Arendt P, Souza L, Oliveira D, et al. 2018. Co-expression of squalene epoxidases with triterpene cyclases boosts production of triterpenoids in plants and yeast. Metabolic Engineering 49: 1–12.

Dufourc EJ. 2008. Sterols and membrane dynamics. Journal of Chemical Biology 1: 63–77.

Espenshade PJ, Hughes AL. 2007. Regulation of Sterol Synthesis in Eukaryotes. Annual Review of Genetics 41: 401–427.

Fabris M, George J, Kuzhiumparambil U, Lawson CA, Jaramillo-Madrid AC, Abbriano RM, Vickers CE, Ralph P. 2020. Extrachromosomal Genetic Engineering of the Marine Diatom Phaeodactylum tricornutum Enables the Heterologous Production of Monoterpenoids. ACS Synthetic Biology 9: 598–612.

Fabris M, Matthijs M, Carbonelle S, Moses T, Pollier J, Dasseville R, Baart GJE, Vyverman W, Goossens A. 2014. Tracking the sterol biosynthesis pathway of the diatom Phaeodactylum tricornutum. New Phytologist 204: 521–535.

Fabris M, Matthijs M, Rombauts S, Vyverman W, Goossens A, Baart GJE. 2012. The metabolic blueprint of Phaeodactylum tricornutum reveals a eukaryotic Entner-Doudoroff glycolytic pathway. The Plant Journal 70: 1004–1014.

Friesen JA, Rodwell VW. 2004. The 3-hydroxy-3-methylglutaryl coenzyme-A (HMG-CoA) reductases. Genome biology 5: 248.

Gallo C, Ippolito G, Nuzzo G, Sardo A, Fontana A. 2017. Autoinhibitory sterol sulfates mediate programmed cell death in a bloom-forming marine diatom. Nature Communications 8: 1–11.

Garaiova M, Zambojova V, Simova Z, Griac P, Hapala I. 2014. Squalene epoxidase as a target for manipulation of squalene levels in the yeast Saccharomyces cerevisiae. FEMS yeast research 14: 310–323.

George J, Kahlke T, Abbriano RM, Kuzhiumparambil U, Ralp PJ, Fabris M. 2020. Metabolic engineering strategies in diatoms reveal unique phenotypes and genetic configurations with implications for algal genetics and synthetic biology. Frontiers in Bioengineering and Biotechnology 8: 513.

Gill S, Stevenson J, Kristiana I, Brown AJ. 2011. Cholesterol-dependent degradation of squalene monooxygenase, a control point in cholesterol synthesis beyond HMG-CoA reductase. Cell Metabolism 13: 260–273.

Gohil N, Bhattacharjee G, Khambhati K, Braddick D. 2019. Engineering strategies in microorganisms for the enhanced production of squalene: advances, challenges and opportunities. Frontiers in Bioengineering and Biotechnology 7: 50.

Gold DA, Caron A, Fournier GP, Summons RE. 2017. Paleoproterozoic sterol biosynthesis and the rise of oxygen. Nature 543: 420–423.

Guillard RRL, Hargraves PE. 1993. Stichochrysis immobilis is a diatom, not a chrysophyte. Phycologia 32: 234–236.

Harker M, Holmberg N, Clayton JC, Gibbard CL, Wallace AD, Rawlins S, Hellyer SA, Lanot A, Safford R. 2003. Enhancement of seed phytosterol levels by expression of an N- terminal truncated Hevea brasiliensis (rubber tree) 3-hydroxy-3-methylglutaryl-CoA reductase. Plant Biotechnology Journal 1: 113–121.

Hildebrand M, Davis AK, Smith SR, Traller JC, Abbriano R. 2012. The place of diatoms in the biofuels industry. Biofuels 3: 221–240.

Jaramillo-Madrid AC, Ashworth J, Fabris M, Ralph PJ. 2019. Phytosterol biosynthesis and production by diatoms (Bacillariophyceae). Phytochemistry 163: 46–57.

Jaramillo-Madrid AC, Ashworth J, Fabris M, Ralph PJ. 2020a. The unique sterol biosynthesis pathway of three model diatoms consists of a conserved core and diversified endpoints. Algal Research 48: 101902.

Jaramillo-Madrid AC, Ashworth J, Ralph PJ. 2020b. Levels of Diatom Minor Sterols Respond to Changes in Temperature and Salinity. Journal of Marine Science and Engineering 8: 85.

Johnson LK, Alexander H, Brown CT. 2019. Re-assembly, quality evaluation, and annotation of 678 microbial eukaryotic reference transcriptomes. GigaScience 8: giy158.

Jones P, Binns D, Chang H-Y, Fraser M, Li W, McAnulla C, McWilliam H, Maslen J, Mitchell A, Nuka G. 2014. InterProScan 5: genome-scale protein function classification. Bioinformatics 30: 1236–1240.

Kajikawa M, Kinohira S, Ando A, Shimoyama M, Kato M, Fukuzawa H. 2015. Accumulation of squalene in a microalga Chlamydomonas reinhardtii by genetic modification of squalene synthase and squalene epoxidase genes. PLoS One 10: e0120446.

Keeling PJ, Burki F, Wilcox HM, Allam B, Allen EE, Amaral-Zettler LA, Armbrust EV, Archibald JM, Bharti AK, Bell CJ. 2014. The Marine Microbial Eukaryote Transcriptome Sequencing Project (MMETSP): illuminating the functional diversity of eukaryotic life in the oceans through transcriptome sequencing. PLoS biology 12: e1001889.

Kim Y, Lee OR, Oh JY, Jang M, Yang D. 2014. Functional analysis of 3-Hydroxy-3- methylglutaryl Coenzyme A reductase encoding genes in triterpene saponin-producing Ginseng. Plant physiology 165: 373–387.

Krogh A, Larsson B, Von Heijne G, Sonnhammer EL. 2001. Predicting transmembrane protein topology with a hidden Markov model: application to complete genomes. Journal of molecular biology 305: 567–580.

Kroth PG. 2002. Protein transport into secondary plastids and the evolution of primary and secondary plastids. In: International Review of Cytology. Academic Press, 191–255.

Kuwabara PE, Labouesse M. 2002. The sterol-sensing domain: multiple families, a unique role? TRENDS in Genetics 18: 193–201.

Lange I, Poirier BC, Herron BK, Lange BM. 2015. Comprehensive assessment of transcriptional regulation facilitates metabolic engineering of isoprenoid accumulation in Arabidopsis. Plant Physiology 169: 1595–1606.

Lange BM, Rujan T, Martin W, Croteau R. 2000. Isoprenoid biosynthesis: the evolution of two ancient and distinct pathways across genomes. Proceedings of the National Academy of Sciences 97: 13172–13177.

Laranjeira S, Amorim-Silva V, Esteban A, Arró M, Ferrer A, Tavares RM, Botella MA, Rosado A, Azevedo H. 2015. Arabidopsis Squalene Epoxidase 3 (SQE3) Complements SQE1 and Is Important for Embryo Development and Bulk Squalene Epoxidase Activity. Molecular Plant 8: 1090–1102.

Leber R, Landl K, Zinser E, Ahorn H, Spo A, Kohlwein SD, Turnowsky F. 1998. Dual localization of squalene epoxidase, Erg1p, in yeast reflects a relationship between the endoplasmic reticulum and lipid particles. Molecular Biology of the Cell 9: 375–386.

Lee M, Jeong J, Seo J, Shin C, Kim Y, In J, Yang D, Yi J, Choi Y. 2004. Enhanced triterpene and phytosterol biosynthesis in Panax ginseng overexpressing squalene synthase gene. Plant Cell Physiology 45: 976–984.

Lee A-R, Kwon M, Kang M-K, Kim J, Kim S-U, Ro D-K. 2019. Increased sesqui- and triterpene production by co-expression of HMG-CoA reductase and biotin carboxyl carrier protein in tobacco (Nicotiana benthamiana). Metabolic Engineering 52: 20–28.

Leivar P, González VM, Castel S, Trelease RN, López-Iglesias C, Arró M, Boronat A, Campos N, Ferrer A, Fernàndez-Busquets X. 2005. Subcellular localization of Arabidopsis 3-hydroxy-3-methylglutaryl-coenzyme A reductase. Plant physiology 137: 57–69.

Li W, Liu W, Wei H, He Q, Chen J, Zhang B, Zhu S. 2014. Species-specific expansion and molecular evolution of the 3-hydroxy-3-methylglutaryl coenzyme A reductase (HMGR) gene family in plants. PLoS One 9: e94127.

Liu X, Hempel F, Stork S, Bolte K, Moog D, Heimerl T, Maier UG, Zauner S. 2016. Addressing various compartments of the diatom model organism Phaeodactylum tricornutum via sub-cellular marker proteins. Algal Research 20: 249–257.

Lohr M, Schwender J, Polle JE. 2012. Isoprenoid biosynthesis in eukaryotic phototrophs: a spotlight on algae. Plant Science 185: 9–22.

Massé G, Belt ST, Rowland SJ, Rohmer M. 2004. Isoprenoid biosynthesis in the diatoms Rhizosolenia setigera (Brightwell) and Haslea ostrearia (Simonsen). Proceedings of the National Academy of Sciences of the United States of America 101: 4413–4418.

Nagai T, Ibata K, Park ES, Kubota M, Mikoshiba K, Miyawaki A. 2002a. A variant of yellow fluorescent protein with fast and efficient maturation for cell-biological applications. Nature Biotechnology 20: 87–90.

Nagai M, Sakakibara J, Nakamura Y, Gejyo F, Ono T. 2002b. SREBP-2 and NF-Y are involved in the transcriptional regulation of squalene epoxidase. Biochemical and Biophysical Research Communications 295: 74–80.

Polakowski T, Stahl U, Lang C. 1998. Overexpression of a cytosolic hydroxymethylglutaryl-CoA reductase leads to squalene accumulation in yeast. Applied Microbiology and Biotechnology 49: 66–71.

Pollak B, Cerda A, Delmans M, Álamos S, Moyano T, West A, Gutiérrez RA, Patron NJ, Federici F, Haseloff J. 2019. Loop assembly: a simple and open system for recursive fabrication of DNA circuits. New Phytologist 222: 628–640.

Pollier J, Vancaester E, Kuzhiumparambil U, Vickers C, Vandepoele K, Goossens A, Fabris M. 2019. A widespread alternative squalene epoxidase participates in eukaryote steroid biosynthesis. Nature Microbiology 4: 226–233.

Rampen SW, Abbas BA, Schouten S, Damste JSS. 2010. A comprehensive study of sterols in marine diatoms (Bacillariophyta): Implications for their use as tracers for diatom productivity. Limnology and Oceanography 55: 91–105.

Ras RT, Geleijnse JM, Trautwein EA. 2014. LDL-cholesterol-lowering effect of plant sterols and stanols across different dose ranges: a meta-analysis of randomised controlled studies. British Journal of Nutrition 112: 214–219.

Re EB, Jones D, Learned RM. 1995. Co-expression of native and introduced genes reveals cryptic regulation of HMG CoA reductase expression in Arabidopsis. The Plant Journal 7: 771–784.

Sasso S, Pohnert G, Lohr M, Mittag M, Hertweck C. 2012. Microalgae in the postgenomic era: a blooming reservoir for new natural products. FEMS Microbiology Reviews 36: 761–785.

Schaller H, Grausem B, Benveniste P, Chye ML, Tan CT, Song YH, Chua NH. 1995. Expression of the Hevea brasiliensis (H.B.K.) Müll. Arg. 3-hydroxy-3-methylglutaryl- coenzyme a reductase 1 in tobacco results in sterol overproduction. Plant Physiology 109: 761–770.

Schreiber U. 2004. Pulse-amplitude-modulation (PAM) fluorometry and saturation pulse method: an overview. In: Chlorophyll a fluorescence. Springer, 279–319.

Sheppard V, Poulsen N, Kröger N. 2010. Characterization of an endoplasmic reticulum- associated silaffin kinase from the diatom Thalassiosira pseudonana. Journal of Biological Chemistry 285: 1166–1176.

Strand TA, Lale R, Degnes KF, Lando M, Valla S. 2014. A new and improved host-independent plasmid system for RK2-based conjugal transfer. PLoS One 9: 1–6.

Theesfeld CL, Pourmand D, Davis T, Garza RM, Hampton RY. 2011. The sterol-sensing domain (SSD) directly mediates signal-regulated endoplasmic reticulum-associated degradation (ERAD) of 3-hydroxy-3-methylglutaryl (HMG)-CoA reductase isozyme Hmg2. The Journal of biological chemistry 286: 26298–26307.

Valitova JN, Sulkarnayeva AG, Minibayeva F V. 2016. Plant sterols: Diversity, biosynthesis, and physiological functions. Biochemistry 81: 819–834.

Vavitsas K, Fabris M, Vickers CE. 2018. Terpenoid Metabolic Engineering in Photosynthetic Microorganisms. Genes 9: 2520.

Vranova E, Coman D, Gruissem W. 2013. Network Analysis of the MVA and MEP Pathways for Isoprenoid Synthesis. Annual review of plant biology 64: 665–700.

Wang J, Huang M, Yang J, Ma X, Zheng S, Deng S, Huang Y, Yang X. 2017. Anti- diabetic activity of stigmasterol from soybean oil by targeting the GLUT4 glucose transporter. Food & Nutrition Research 61: 1364117.

Zhang D-H, Jiang L-X, Li N, Yu X, Zhao P, Li T, Xu J-W. 2017. Overexpression of the squalene epoxidase gene alone and in combination with the 3-Hydroxy-3-methylglutaryl Coenzyme A gene increases ganoderic acid production in Ganoderma lingzhi. Journal of Agricultural and Food Chemistry 65: 4683–4690.

