## Supplementary material for "Overexpression of key sterol pathway enzymes in two model marine diatoms alters sterol profiles in *Phaeodactylum tricornutum*"

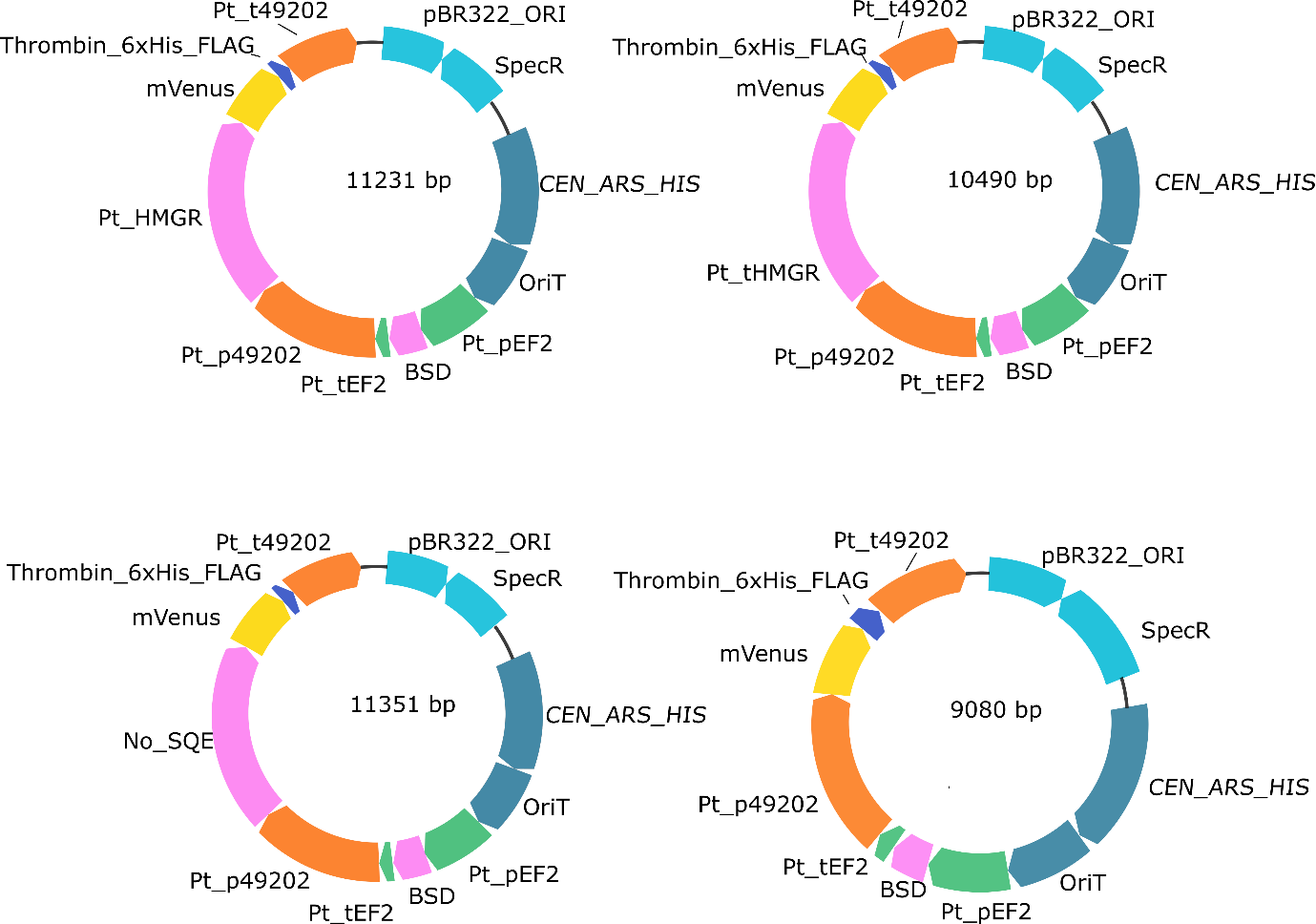

Figure S1. Maps of plasmids used for *P. tricornutum* transformation. CDS are shown in pink, in green promoters and terminators and fluorescence mVenus gene in yellow.

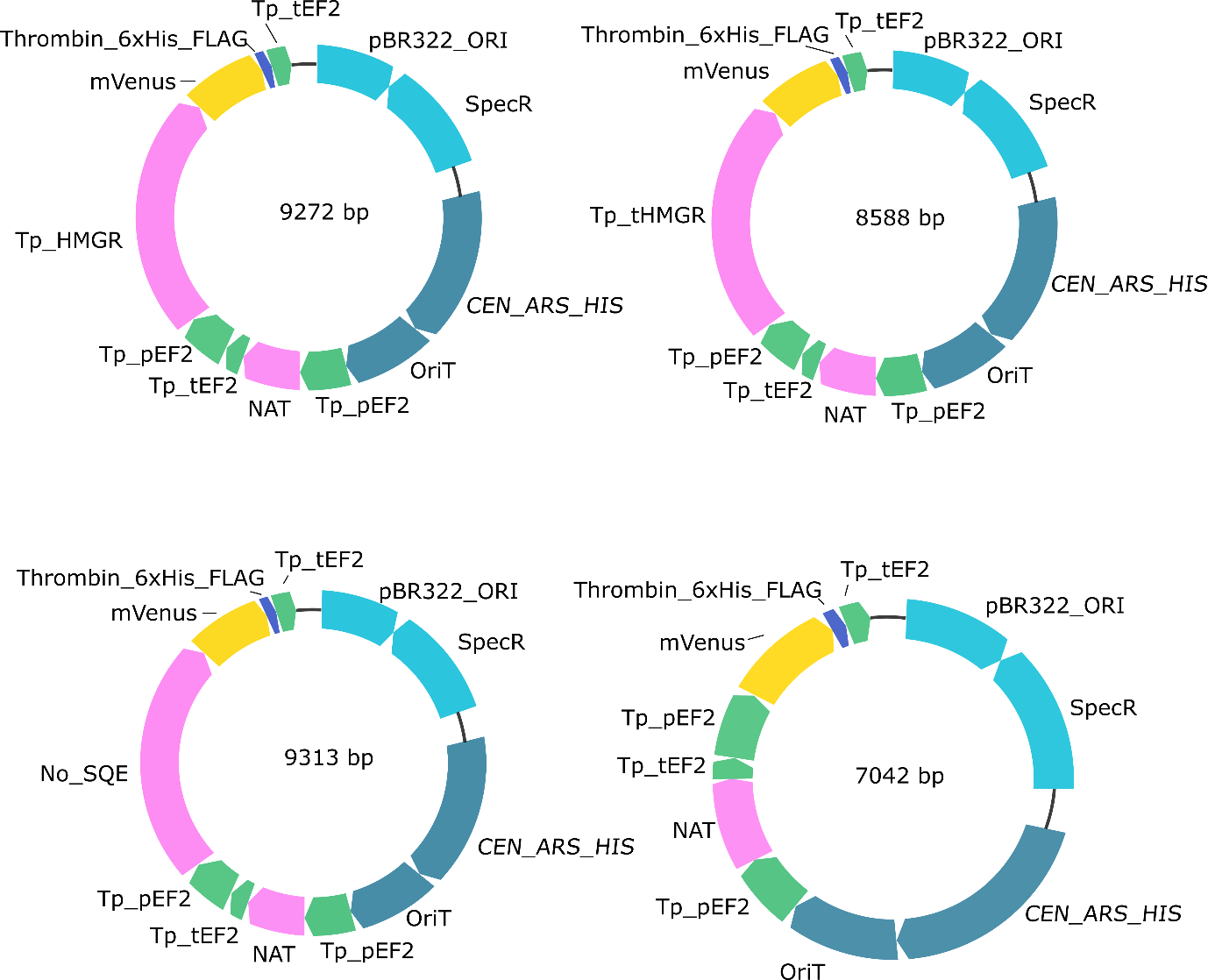

Figure S2. Maps of plasmids used for *T. pseudonana* transformation. CDS are shown in pink, in green promoters and terminators and fluorescence mVenus gene in yellow.

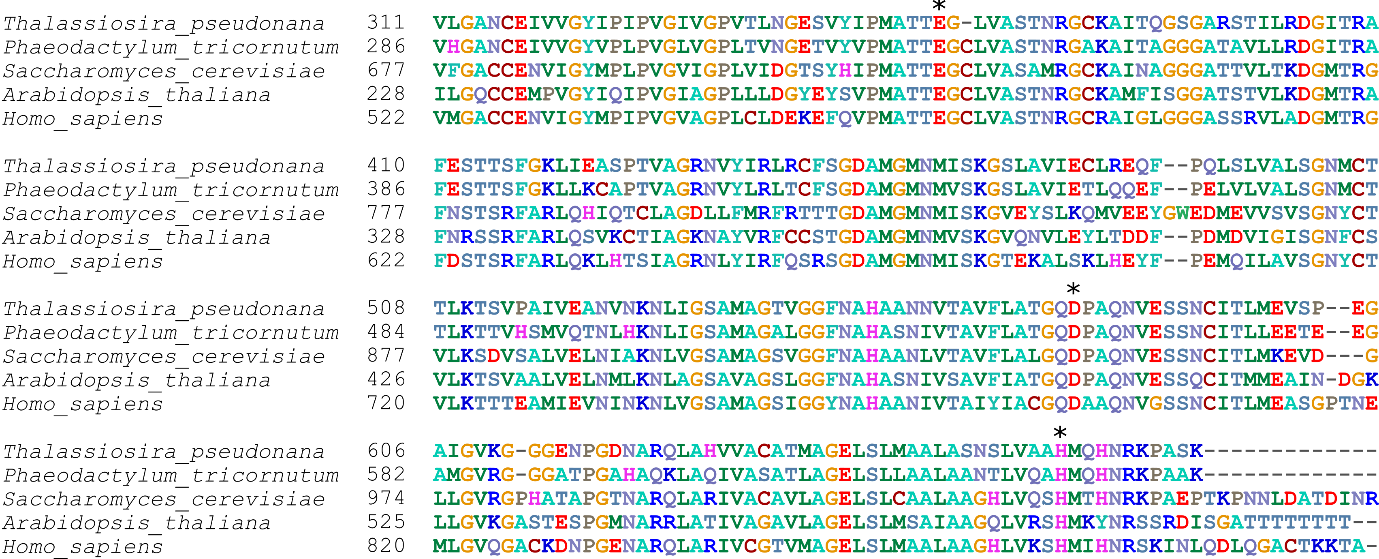

Figure S3. Sequence alignment of HMGR protein (catalytic domain) from model organisms. Asterisks (*) indicate conserved catalytic residues.

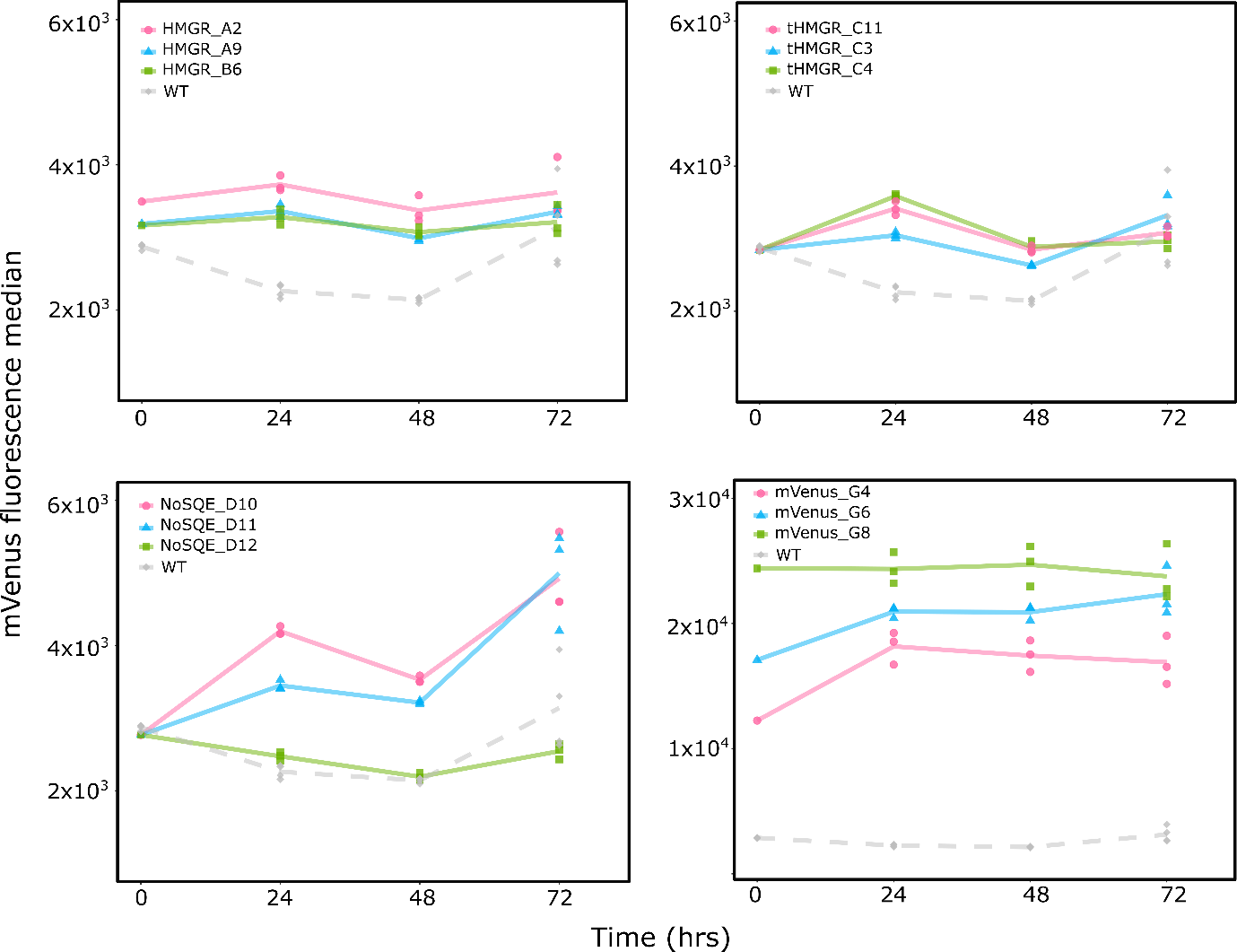

Figure S4. mVenus fluorescence during full scale experiment in *P. tricornutum* transformants (n = 3).

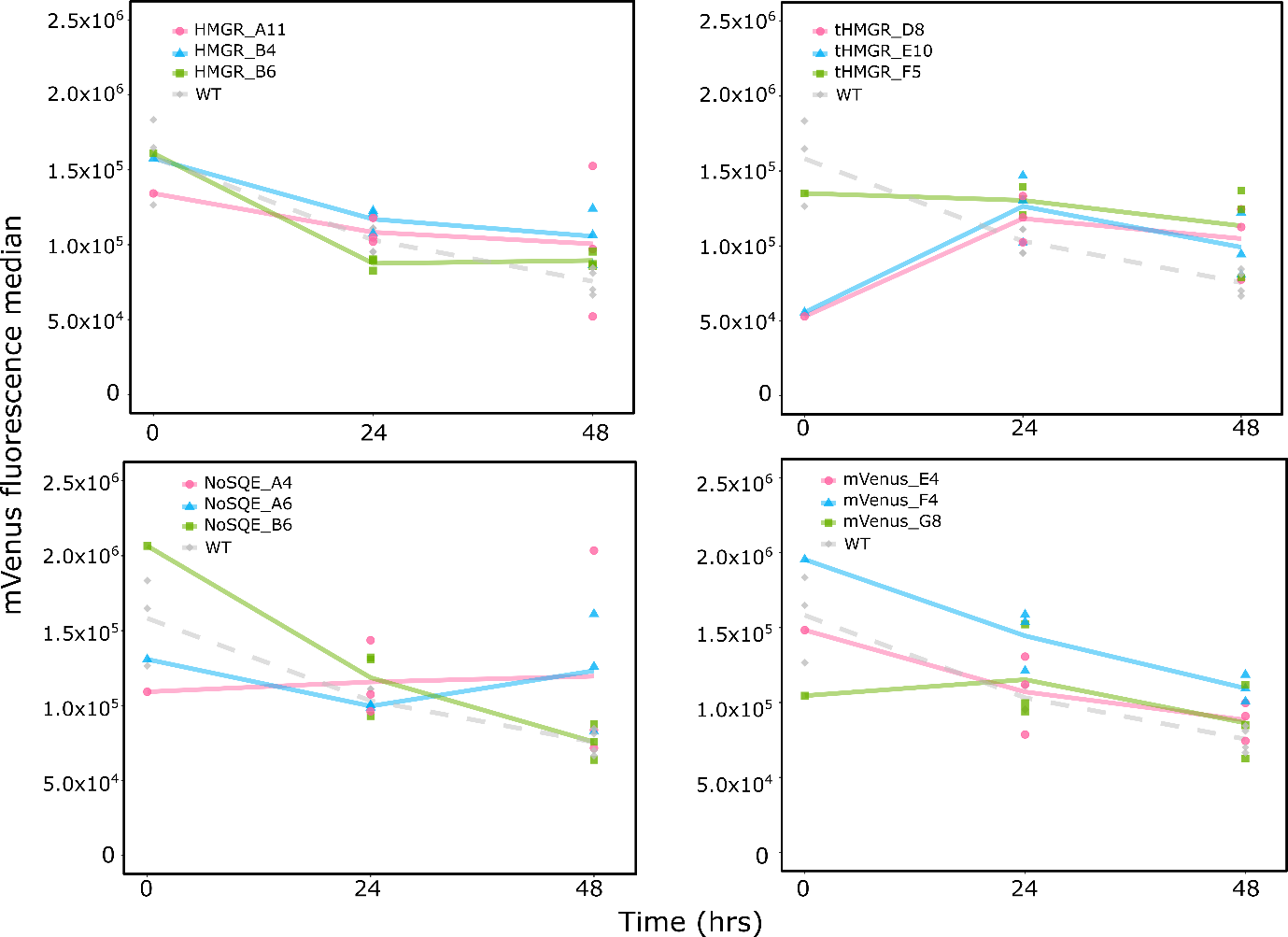

Figure S5. mVenus fluorescence during full scale experiment in *T. pseudonana* transformants (n = 3).

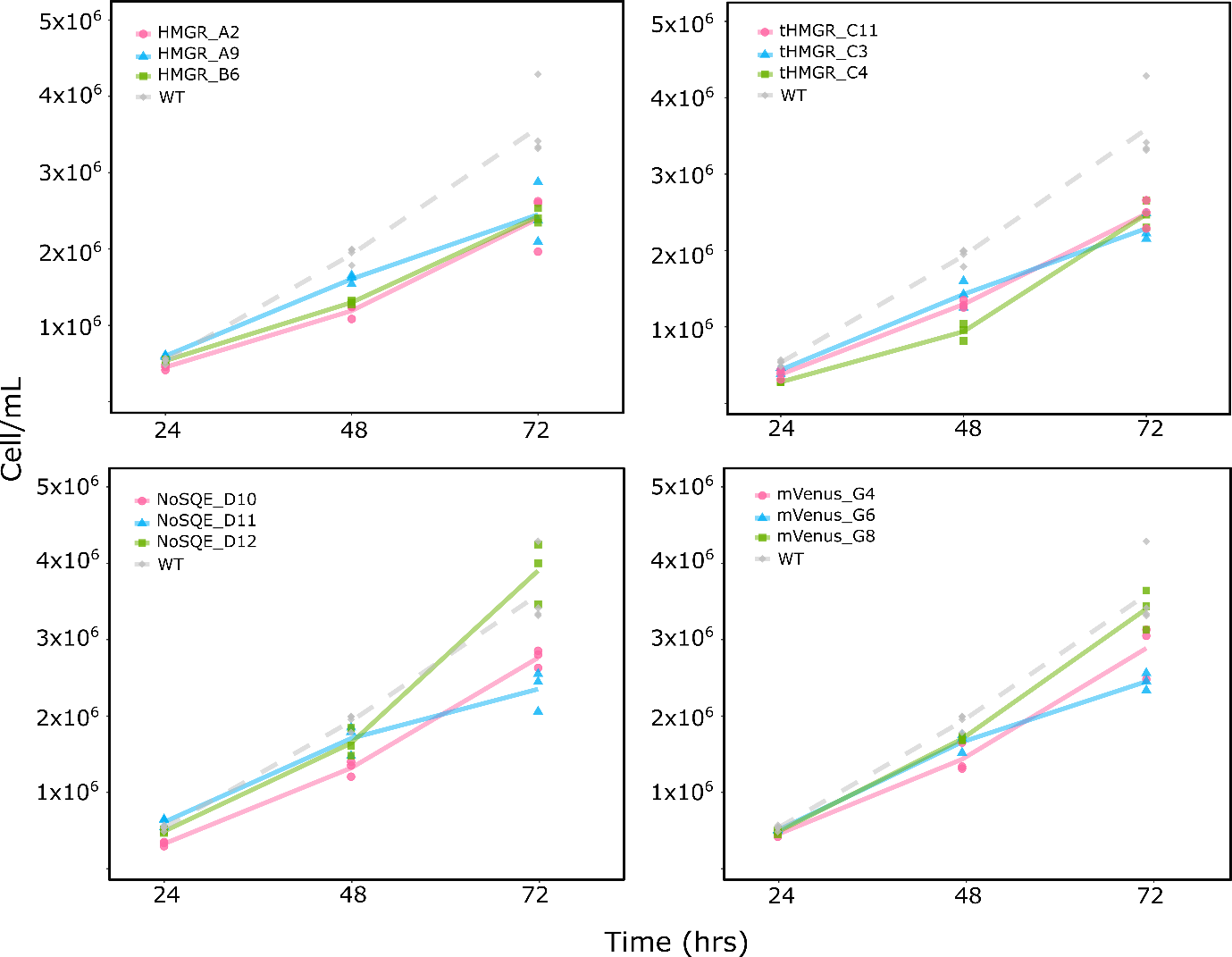

Figure S6. Growth curves during full scale experiment for *P. tricornutum* transformants (n = 3).

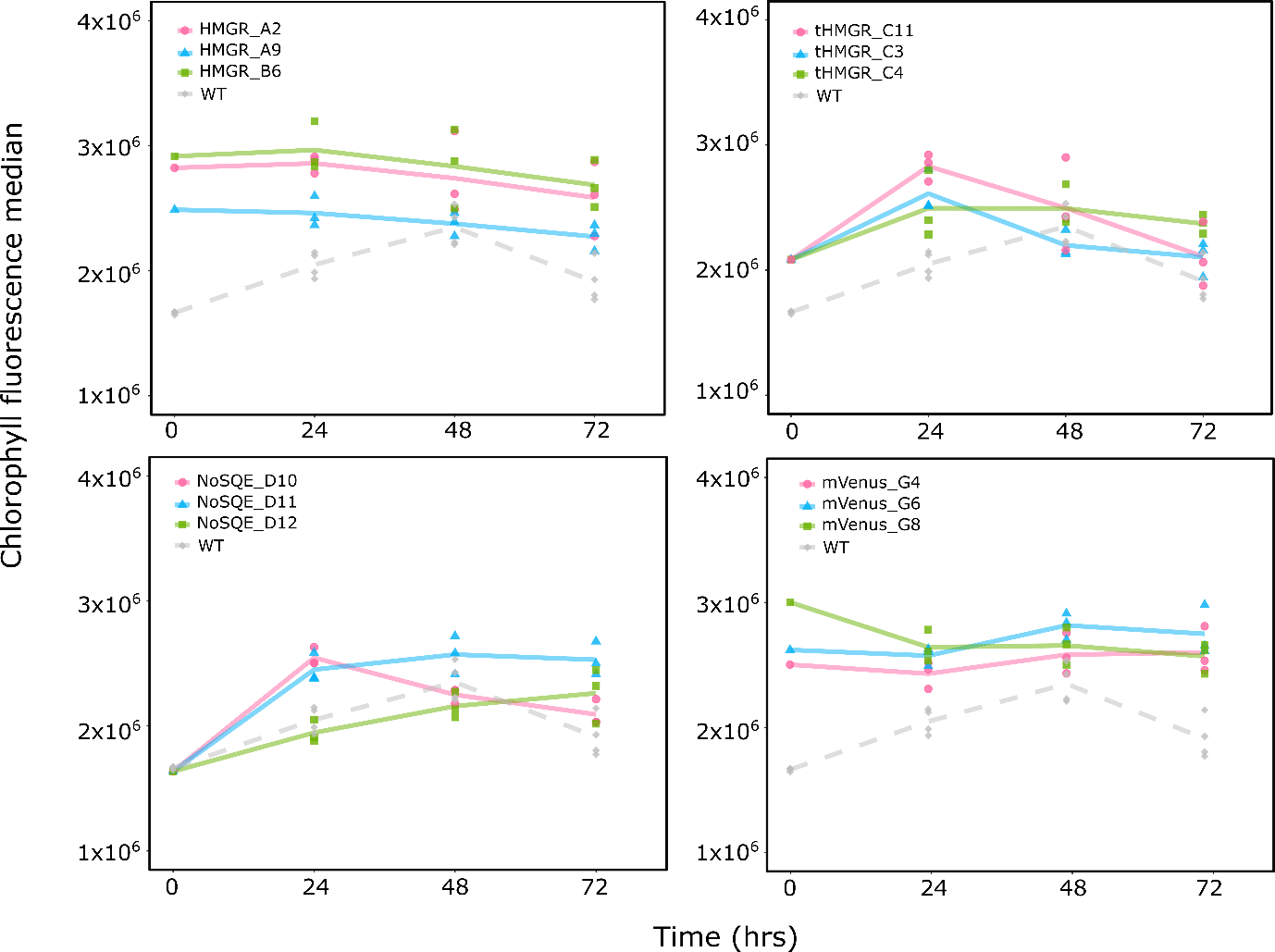

Figure S7. Chlorophyll fluorescence during full scale experiment in *P. tricornutum* transformants (n = 3).

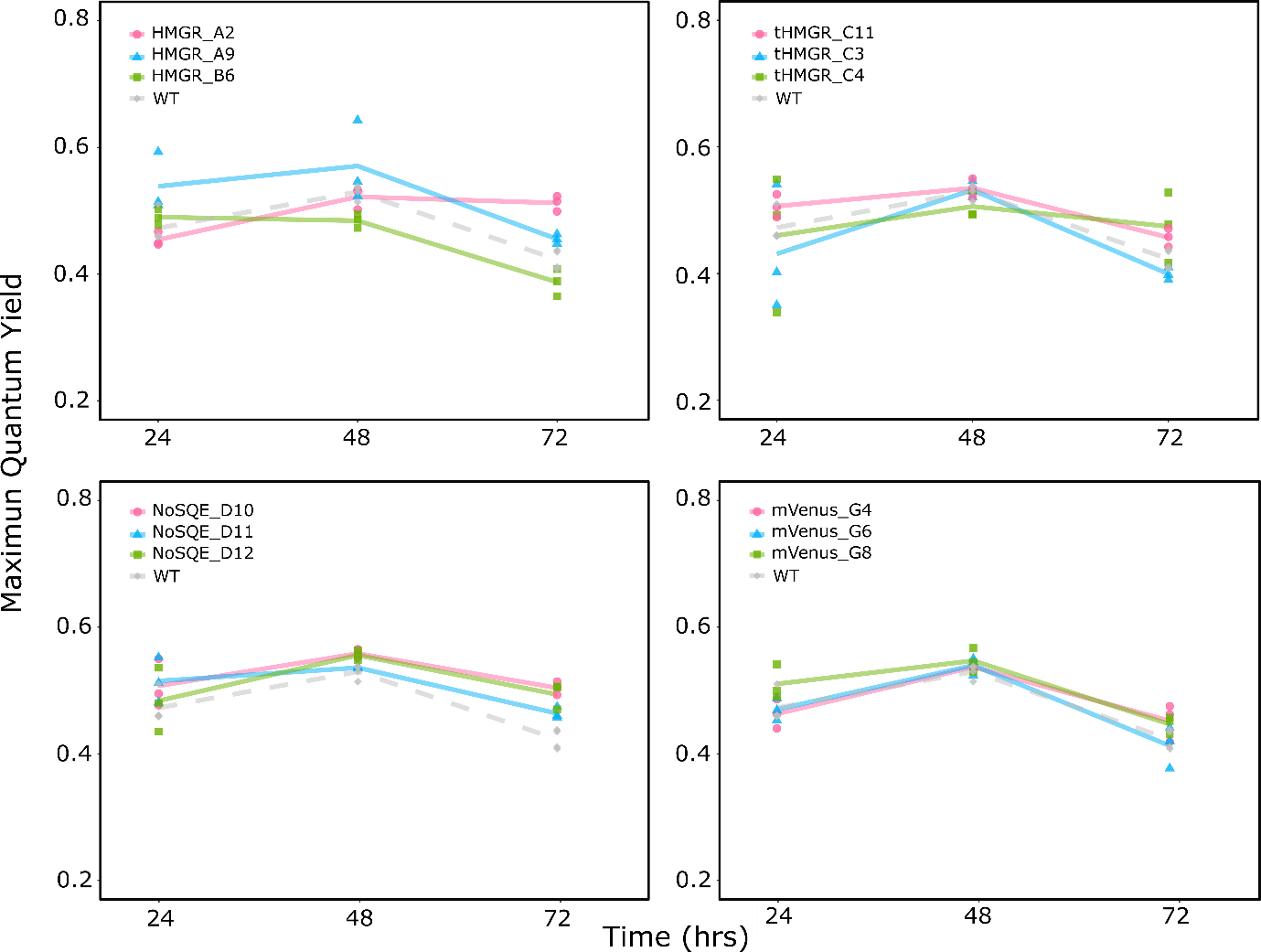

Figure S8. Maximum quantum yield for *P. tricornutum* transformants during full scale experiment (n = 3).

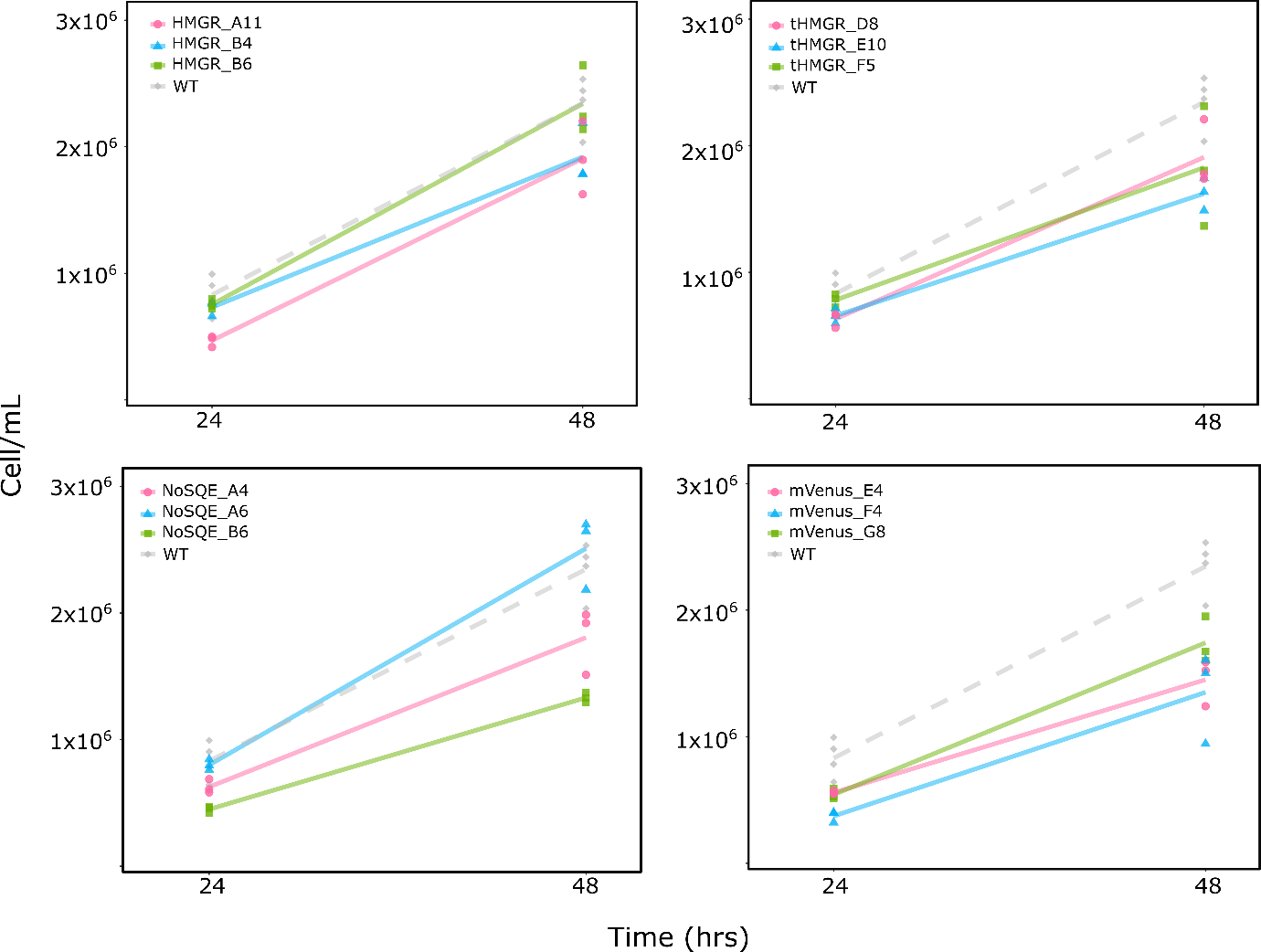

Figure S9. Growth curves during full scale experiment for *T. pseudonana* transformants (n = 3).

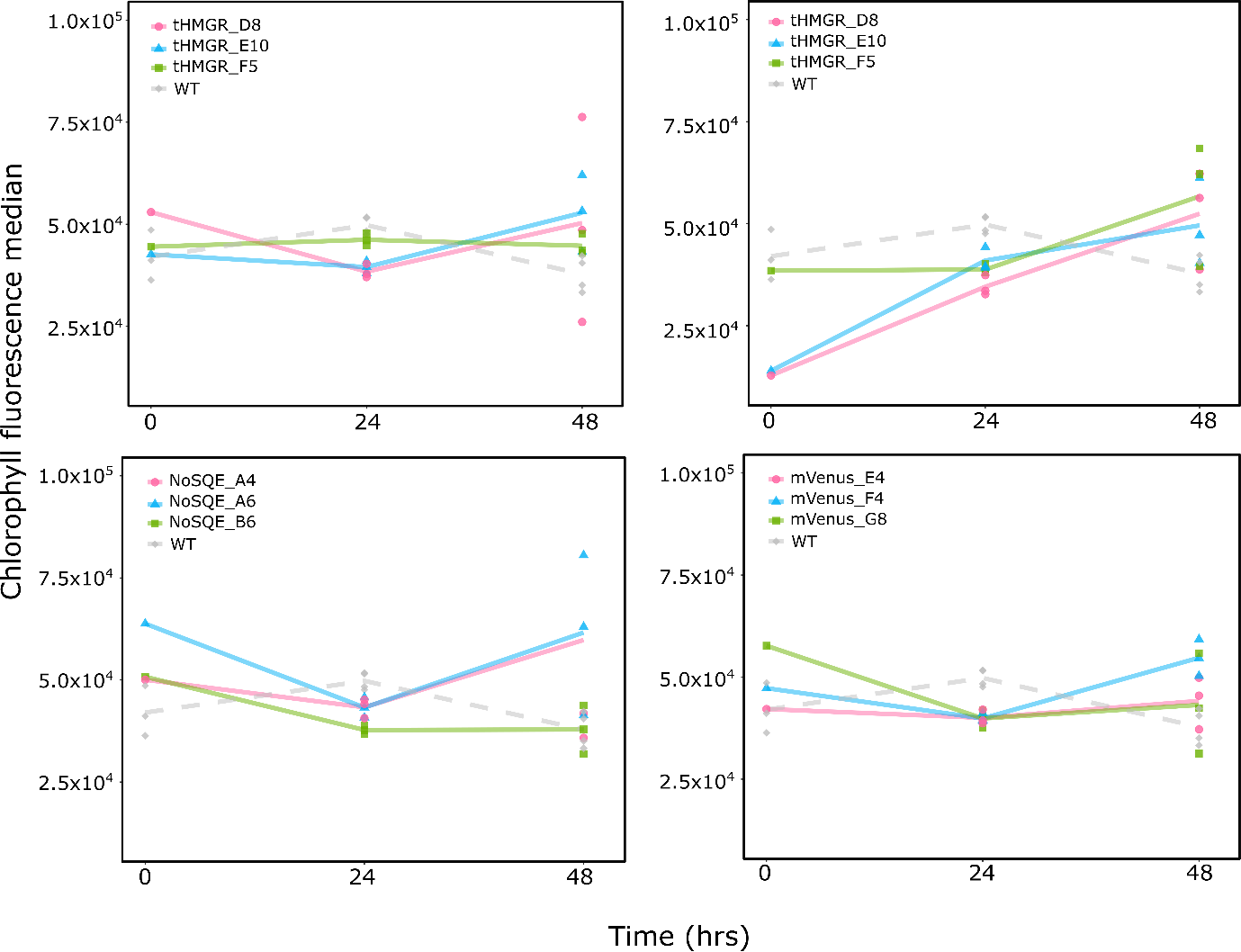

Figure S10. Chlorophyll fluorescence during full scale experiment in *T. pseudonana* transformants (n = 3).

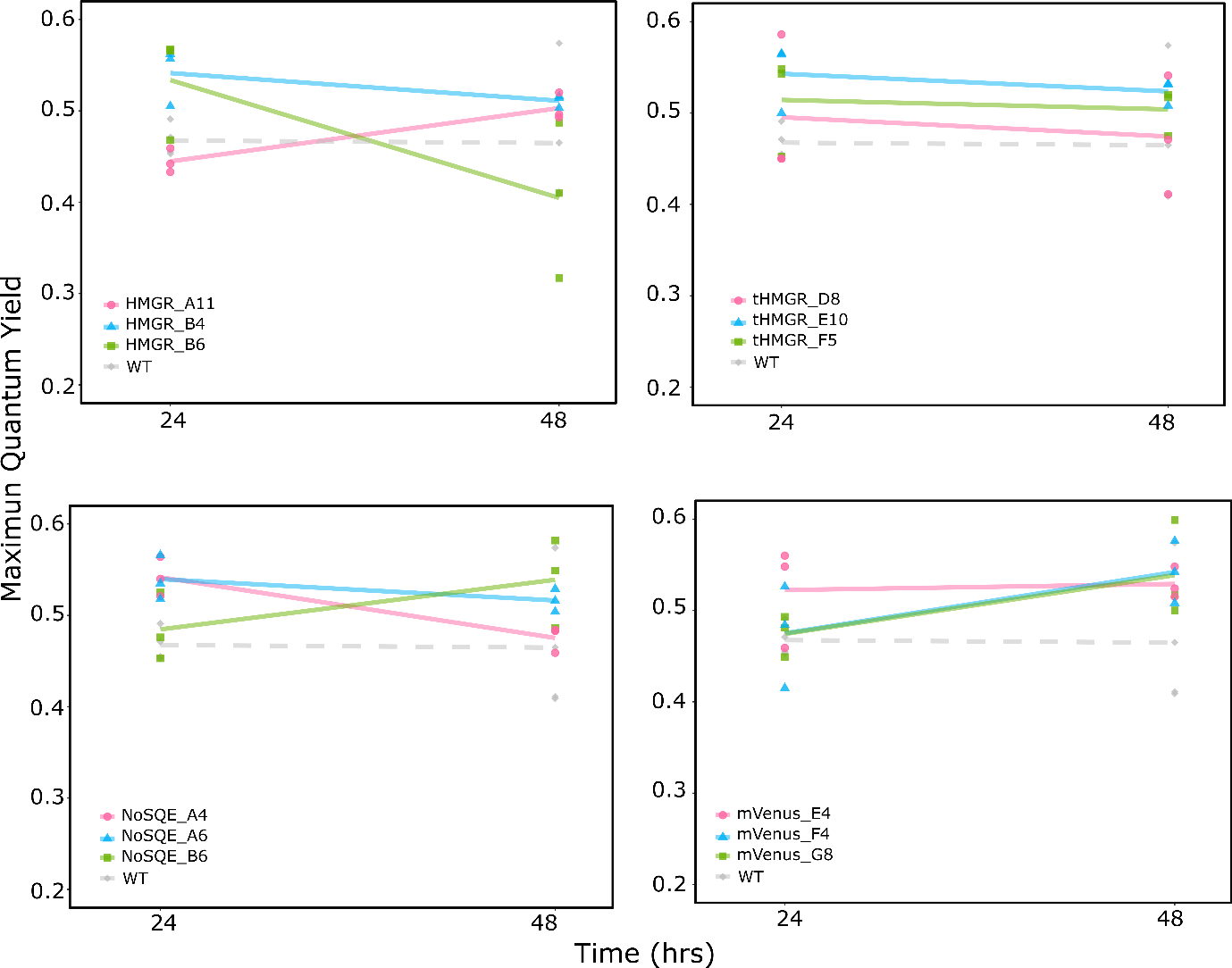

Figure S11. Maximum quantum yield for *T. pseudonana* transformants during full scale experiment (n = 3).

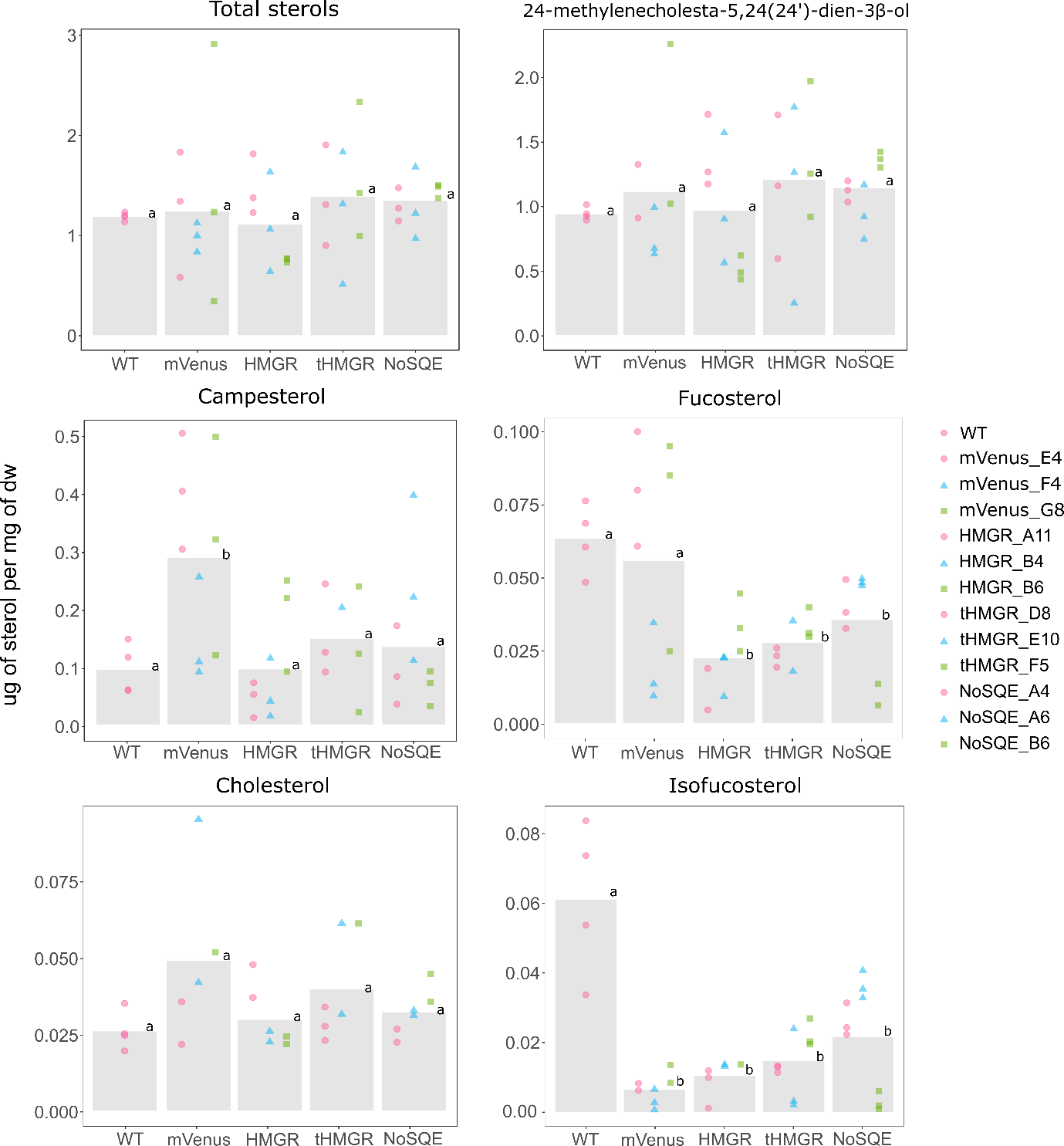

Figure S12. Sterol levels in *T. pseudonana* transformants. Identical letters denote no statistically significant differences among groups using the Pairwise Wilcoxon Rank Sum tests (*p* < 0.05, n = 9).

| **Table S1** L0 parts for construction of episomes using uLoop assembly method (Pollak et al., 2018). Primers sequences used for domestication are presented for L0 each part | | | | | |
| --- | --- | --- | --- | --- | --- |
| **L0 part** | **Gene ID** | **Description** | **Template** | **Primers** | **Plasmid number** |
| AC_pTpEF2 | 269148 | Elongation factor promoter | *P. tricornutum* genomic DNA | F: AAGCTCTTCATCCggagAGTGTGCAATGCAGTCAATTCAATAGATATG  R: TTGCTCTTCTTCGcattCTTGACGTTCTTTTCTCTTTAATTAATCGCG |  |
| AC_pPtEF2 | Phatr3_J35766 | Elongation factor promoter | *P. tricornutum* genomic DNA | F: AAGCTCTTCATCCggagTCCATTTTGACATGTTTCCTAGCTAGAAG  R: TTGCTCTTCTTCGcattTGTGTGGAGAGAACGAGCAGCAGCG |  |
| AC_pPt49202 | Phatr3_J49202 | Predicted protein JCVI | NA | NA |  |
| AC_CENARSHIS | NA | CEN6-ARSH4-HIS3 episome maintenance sequence from PtPBR11 (Genebank KX523203) (Diner *et al.*, 2016) | NA | NA |  |
| CF_OriT | NA | Origin of transfer, originally called basis of mobilization (Diner *et al.*, 2016) | PtPBR11 (Genebank KX523203) | F: CAGAAGCTCTTCATCCaatgGATCGTCTTGCCTTGCTCGTCGG  R: TTGCTCTTCTTCGagcgATCTTCCGCTGCATAACCCTGCTTCGG | 139948 |
| CD_NoSQE | 521007 | 0.0 | NA | NA |  |
| CD_TpHMGR | 33680 | 3-hydroxy-3-methylglutaryl-coenzyme A reductase (EC: 1.1.1.34) | *T. pseudonana* genomic DNA | F: AAGCTCTTCATCCaatgGCGGCAGCAGCAGCACCAACCATAG  R: TTGCTCTTCTTCGacctgaCTTTGAAGCAGGCTTGCGATTATGTTG |  |
| CD_TptHMGR | 33680 | Truncated 3-hydroxy-3-methylglutaryl-coenzyme A reductase (EC: 1.1.1.34) | *T. pseudonana* genomic DNA | F: AAGCTCTTCATCCaatgAGCCCCAACGACCCACCCGTCAAAG  R: TTGCTCTTCTTCGacctgaCTTTGAAGCAGGCTTGCGATTATGTTG |  |
| CD_PtHMGR | Phatr3_J16870 | 3-hydroxy-3-methylglutaryl-coenzyme A reductase | *P. tricornutum* genomic DNA | F: AAGCTCTTCATCCaatgACGGTGACTATCAGCAGTAGTATTAG  R: TTGCTCTTCTTCGacctgaCTTGGCGGCGGGTTTGCGGTTG |  |
| CD_PttHMGR | Phatr3_J16870 | Truncated 3-hydroxy-3-methylglutaryl-coenzyme A reductase | *P. tricornutum* genomic DNA | F: AAGCTCTTCATCCaatgGACTCTATTTCCACCAAGACCAGCGCG  R: TTGCTCTTCTTCGacctgaCTTGGCGGCGGGTTTGCGGTTG |  |
| CD_bsd | NA | blasticidin-S deaminase | pST1374-N-NLS-flag-linker-Cas9-D10A (Addgene #51130) | F: AAGCTCTTCATCCaatgGCCAAGCCTTTGTCTCAAGAAGAATCCAC  R:TTGCTCTTCTTCGacctgaGCCCTCCCACACATAACCAGAGGG |  |
| CD_nat | NA | nourseothricin acetyltransferase | pAGM4723:TpCC_Urease (Addgene #85982) | F: AAGCTCTTCATCCaatgACCACTCTTGACGACACGGCTTACCGG  R:TTGCTCTTCTTCGacctgaGGGGCAGGGCATGCTCATGTAGAGCG |  |
| CD_mVenus |  | Yellow fluorescent protein | NA | NA |  |
| DE_Thr6XHisFLAG | NA | General C-terminal protein tag. Thormbine, 6xHIS and FLAG (DYKDDDDK) tag | DE_Venus-ThrHISFLAG L0 part | F: AAGCTCTTCATCCaggtGCGCTGGTCCCTCGCGGTAG  R: TTGCTCTTCTTCGaagcTCACTTATCGTCATCATCCTTGTAGTCG |  |
| DE_Venus-ThrHISFLAG | NA | Yellow fluorescent protein with a Thormbine, 6xHIS and FLAG (DYKDDDDK) tag JCVI | NA | NA |  |
| DE-3xStop | NA | Three stop codon | NA | NA |  |
| EF_tPtEF2 | Phatr3_J35766 | Elongation factor terminator | *P. tricornutum* genomic DNA | F: AAGCTCTTCATCCGCTTACAGAAAAACAGACTCATAGGGTAC  R:TTGCTCTTCTTCGAGCGGAGACATTACTCCACACGATGG |  |
| EF_tTpEF2 | 269148 | Elongation factor terminator | *T. pseudonana* genomic DNA | F: AAGCTCTTCATCCgcttATATCTTCTTGCAACAATGGTAGCG  R: TTGCTCTTCTTCGagcgAGCAGGGTTGGTTAGAGATAACTAATG |  |
| EF_tPt49202 | Phatr3_J49202 | Predicted protein | NA | NA |  |

**Table S2**. Source and number of transmembrane domains of *HMGR* sequences used for phylogenetic and domain analysis

| **Organism** | **MMETSP ID** | **Present** | **Source** | **Strain** | **Gene ID** | **TM*** |
| --- | --- | --- | --- | --- | --- | --- |
| *Attheya septentrionalis* | MMETSP1449 | Y | Transcriptomics | CCMP2084 | Transcript_1390 | 3 |
| *Chaetoceros affinis* | MMETSP0091 | Y | Transcriptomics | CCMP159 | Transcript_17836 | 0 |
| *Chaetoceros brevis* | MMETSP1435 | N | Transcriptomics | CCMP164 | NA | NA |
| *Chaetoceros curvisetus* | MMETSP0716 | N | Transcriptomics | Unknown | NA | NA |
| *Chaetoceros debilis* | MMETSP0149 | N | Transcriptomics | MM31A-1 | NA | NA |
| *Chaetoceros dichaeta* | MMETSP1447 | Y | Transcriptomics | CCMP1751 | Transcript_4115 | 0 |
| *Chaetoceros muelleri* | NA | N | Transcriptomics | CCMP1316 | NA | NA |
| *Chaetoceros neogracile* | MMETSP0752 | Y | Transcriptomics | CCMP1317 | Transcript_689 | 3 |
| *Chaetoceros sp.* | MMETSP1429 | Y | Transcriptomics | UNC1202 | Transcript_18209 | 0 |
| *Coscinodiscus wailesii* | MMETSP1066 | Y | Transcriptomics | CCMP2513 | Transcript_2372 | 3 |
| *Cylindrotheca closterium* | MMETSP0017 | Y | Transcriptomics | KMMCC:B-181 | Transcript_33356 | 3 |
| *Ditylum brightwelli* | MMETSP1001 | Y | Transcriptomics | GSO105 | Transcript_14074 | 3 |
| *Extubocellulus spinifer* | MMETSP0696 | Y | Transcriptomics | CCMP396 | Transcript_715 | 3 |
| *Fistulifera solaris* | NA | Y | Genome | JPCC DA0580 | GAX29119.1 | 3 |
| *Fragilariopsis kerguelensis* | MMETSP0909 | Y | Transcriptomics | L2-C3 | CAMPEP_0196091342 | 2 |
| *Homo sapiens* | NA | Y | Genome | NA | 3156 | 5 |
| *Leptocylindrus danicus* | MMETSP0321 | Y | Transcriptomics | B650 | Transcript_1399 | 2 |
| *Nitzschia punctate* | MMETSP0747 | Y | Transcriptomics | CCMP561 | Transcript_27671 | 3 |
| *Nitzschia sp* | MMETSP0014 | Y | Transcriptomics | RCC80 | Transcript_11673 | 3 |
| *Odontella Sinensis* | MMETSP0160 | Y | Transcriptomics | Grunow 1884 | Transcript_15934 | 3 |
| *Oryza sativa* | NA | Y | Genome | NA | LOC_Os02g48330 | 0 |
| *Phaeodactylum tricornutum* | NA | Y | Genome | CCMP632 | Phatr3_J16870 | 3 |
| *Pseudonitzschia arenysensis* | MMETSP0329 | Y | Transcriptomics | B593 | CAMPEP_0116128146 | 3 |
| *Pseudonitzschia delicatissima* | MMETSP0327 | Y | Transcriptomics | B596 | CAMPEP_0116102328 | 3 |
| *Pseudonitzschia multiseries* | NA | Y | Genome | CLN-47 | 288249 | 3 |
| *Pseudonitzschia pungens* | MMETSP1061 | Y | Transcriptomics | cf. pungens | Transcript_13267 | 3 |
| *Rhizosolenia setigera* | MMETSP0789 | Y | Transcriptomics | CCMP1694 | CAMPEP_0178949462 | 3 |
| *Saccharomyces cerevisiae* | NA | Y | Genome | ATCC 204508 | 854900 | 7 |
| *Skeletonema marinoi* | MMETSP1428 | Y | Transcriptomics | UNC1201 | Transcript_20411 | 3 |
| *Skeletonema menzelii* | MMETSP0603 | Y | Transcriptomics | CCMP793 | Transcript_7316 | 3 |
| *Solanum lycopersicum* | NA | Y | Genome | NA | Solyc02g082260.3 | 2 |
| *Thalassiosira miniscula* | MMETSP0737 | Y | Transcriptomics | CCMP1093 | Transcript_23358 | 2 |
| *Thalassiosira oceanica* | NA | Y | Genome | CCMP1005 | 91521 | 2 |
| *Thalassiosira pseudonana* | NA | Y | Genome | CCMP1335 | 269148 | 3 |
| *Thalassiosira puntigera* | MMETSP1067 | Y | Transcriptomics | Tpunct2005C2 | Transcript_37597 | 3 |
| *Thalassiosira rotula* | MMETSP0403 | Y | Transcriptomics | CCMP3096 | Transcript_15672 | 2 |
| *Thalassiosira weissflogii* | MMETSP1414 | Y | Transcriptomics | CCMP1010 | Transcript_6235 | 3 |
| *Thalassiothrix antarctica* | MMETSP0152 | Y | Transcriptomics | L6-D1 | Transcript_7954 | 3 |

*TM indicates number of transmembrane domains predicted by TMHMM Server v. 2.0 (http://www.cbs.dtu.dk/services/TMHMM-2.0/). One of the three transmembrane domains in Arabidopsis is located in C-terminus domain.

**HMGR sequences for** **phylogenetic reconstruction**

>Thalassiosira_pseudonana

MAAAAAPTIGQRLDTLLAALSNIGDMNHLLSNYQPSTSQIYSLVIVLSVAFSFRLLNSGDDLGSKLSSSTVEQAKRGEKGTSSKYAKANNKNNKQSYDFPQPKWHILKLTNYVVTTLFLLSIITFLSNASVYLNDNTALMTFLGVWSGLLCYFFGFFGISFVELDDLVVTDADGQQRQQGVVTQPQKQASCKSRKVESPNDPPVKVISLHPPASSTPVCSDPIKSKSPTNTGIPTNVKDLSNEEIATLVLQDKIKDHQLEKLLDPHRAVAVRRLKFDALLSSLGKTTGEDDKKGGVLSELPHEHDLDYKRVLGANCEIVVGYIPIPVGIVGPVTLNGESVYIPMATTEGLVASTNRGCKAITQGSGARSTILRDGITRAPCVRLPSAHEAAQVHLWIEEADNFAKLKEAFESTTSFGKLIEASPTVAGRNVYIRLRCFSGDAMGMNMISKGSLAVIECLREQFPQLSLVALSGNMCTDKKAAAMNWIEGRGKSVVIEATIPKDVVRSTLKTSVPAIVEANVNKNLIGSAMAGTVGGFNAHAANNVTAVFLATGQDPAQNVESSNCITLMEVSPEGDLWISCTMPSIEVGTVGGGTGLSAQSACLRAIGVKGGGENPGDNARQLAHVVACATMAGELSLMAALASNSLVAAHMQHNRKPASK

>Phaeodactylum_tricornutum

MTVTISSSISSSIGSSTTAPTLGMQVDALVQQMDNLSSTQLYGLIVGLTILVSFVLLGSSADIPVSLRDT

TKETSSSSSTPRKTTTVSSSSNRGPEPRWHIFTYVNYAIVACFVASVAEFGRNASAYLAADDNVVLYFLV

AWSVFLCYFFGFFGVSFVHDADAAVASPTPTKPTVRDSISTKTSASNTLSARHPPAPSAPVCSDPSSFTP

IHTSKTNITTLDNAAICQLVLTNQIKDHELEKRLDAHRAVQVRRLVVAHKLDTLEHINAHALDNLPSEPS

LDYTRVHGANCEIVVGYVPLPVGLVGPLTVNGETVYVPMATTEGCLVASTNRGAKAITAGGGATAVLLRD

GITRAPCVRMPSAAQAAHLKLWCETPQHFSTLKRAFESTTSFGKLLKCAPTVAGRNVYLRLTCFSGDAMG

MNMVSKGSLAVIETLQQEFPELVLVALSGNMCTDKKAAATNWLEGRGKSIVVEATIPKDVVTNTLKTTVH

SMVQTNLHKNLIGSAMAGALGGFNAHASNIVTAVFLATGQDPAQNVESSNCITLLEETEEGDLWISCTMPSIEVGTVGGGTSLPAQAACLQAMGVRGGGATPGAHAQKLAQIVASATLAGELSLLAALAANTLVQAHMQHNRKPAAK*

>Fistulifera_solaris

MESLRLQLSQALSSFVSVSSNQLYGVIVGVTVFGCWGILSLAEQPELPEWQKTAVQRVPRQKEGPEPQWHLFRWINMLVVVAFACSVAEFSLNATAYDSNVLCQFLMAWSLLLCYFFGFFGVMFVHDLEDSETVSTPVTKSSKIHPPASSTPICSDAIQSKQNKLLIIEDIKDKKDAEIAQLVFTNQIKDHELEKQLDCHRAVTVRRLVVEQKLASLGHSGALEGLPYETDLDYSRVHGANCEIVVGYVPLPVGIIGPLTINGESFYIPMATTEGCLVASTNRGAKAITEGGGALAQIVRDGITRAPCIRMPTAMAAAQLKLWCETPENFVQMKAAFESTTSFGKLHECKATIAGKNVYLRLVCFSGDAMGMNMVSKGTLAVIECLRKEFPELSLVALSGNMCTDKKAAATNWLEGRGKSVVVEATIPKDVVRKTLKTTVPAIVETNLNKNLVGSAMAGVIGGFNAHASNIVSAIFLATGQDPAQNVESSNCITLMEETESGDLWISCTMPSIEVGTVGGGTSLAAQAACLKALGCQGGGANPGDNAQRLAKVIASATMAGELSLLAALAANTCTYLLTDGFWRTNINSPIPSFFSVVAAHMQHNRKPTK

>Pseudonitzschia_multiseries

MAETMGTMPTLGMRLDNMIAHIDQIPSTQIYGLIVAATVGLCVVLLGTGNSNLEIHHQRQMRLQQQSQNDDLKKPKMAASARSGKQPKWHLFKWINFVAVGGFLYSVFTFCSNASRYLHHESQGVLVQFLVGWSIFLMYFFGFFGVSLIHEDIPREEEESASIPSDPSSFKTSKASSSIPENLKELGNDEIAALVLDNKVKDHMLEKLLD

PFRAVTVRRIACNQKLAAVLGRNNNDTNVLDKLPSEPSLDYSRVFGANCEIVVGYVPLPVGLVGPLTIND

ESVYVPMATTEGCLVASSNRGAKAITQGGGARARIVRDGITRAPCLRMNSAMEAADLKIWCELPQNFAIL

KQAFESTTSFGKLLECNPTVAGKNVYLRLVCFSGDAMGMNMVSKGSLAVIETLQKQFPTCQLIALSGNMC

TDKKAAATNWLLGRGKSVVVECVIPKEVVRTTLKTTVAALVHTNLNKNLIGSAMAGAIGGFNAHASNIVT

AVFLATGQDPAQNVESSNCITLMEEQDDGDLWMCCTMPSIEVGTVGGGTSLPAQAACLEAIGCKGGGSTPGANAKKLATVVAAATMAGELSLLAALAANTLVQAHMVHNRKPAAKK*

>Thalassiosira_oceanica

MMAGTATIGQRLDALVATAQANTDGSAALFALVIGISVSFSFYLLNSGSGSTASAEMTKCGAQLSTCPKD

NRTAIQKPRSPEDPPEPNWNALKLTNCVAATGFLLSVLKFASNASAYLNDSTSLLQFLSIWSVFLLYFFG

FFGIALVDLDGLVEEQPSPAPAAKSRQPISPSKKIAKTVDVEPSPPVKVVNVHPPAKSAPVCSDPSSLKR

HASSVPSNLKEMPNEEIAALVLQDKIKDHQLEKLLDPHRAVEVRRLKFDAKLKSLGNGGALNDLPHKHDL

DYKRVLGANCEIVVGYVPIPVGLAGPITLNGESVYIPMATTEGCLVASTNRGCKAITQGSGAFSTILKDG

ITRAPCCRLPSAQQAAEVAIWIETPENFAELKRAFESTTSFGKLIEARPTVAGRNVYIRLKCFAGDAMGM

NMISKGSLAVIECLRSIFPALSLLALSGNVCTDKKAAAMNWIEGRGKSVVIEATIPKDVVSKTLKTSVKA

IVDANVNKNLIGSAMASVVGGFNAHAANNVTAVFLATGQDPAQNVESSNCITIMEETSDGNLWVSCTMPSIEVGTVGGGTGLPAQAACLRAIGVKGGGEKPGQNAKQLAHVVAAATLAGELSLMAALASNSLVAAHMAHNRKPPTASKVEKAFDEVQHGFDDVWLSGQQQHGKKLLDRCSHPHKNWLLGLLIKLQARLRSRLTVAVGLAPRPQDHRYFIGITRYAPLGPIIGISSALRVMRLWDQSRPKQIWQRFLTDPTQTPYTIHDQLLGLTLVLKSAKNLQKSEMTPAAERRGSINRGYRITDSTCGFCSTEVCEWITDRLDHDNWDTCSGQRQGASWTNNTAIQSLVVGGQEPARAPLYLHYVPRLYKAHARSKITGVVCFRLQIDPLSYASWR

>Extubocellulus_spinifer

MTTVPTLGERLDALVDRIDSIPSTQLYIAAVVTSVVFSCMLLNSGGSNGVVDPAKIPDNI

GRPPSVRRKRRDGSNPEDEPEPRWHILTILNYAAVTSFTSSVLYFATDASRFLADSSSLL

KFITGWSVFLCYFFGFFGISFVDADGLIAREEHEAAARQQQQRMQKQHSLGISSPPSKSK

SKLHSAACSVPVSSDGASTKKRSSSSSCTPVDVKELADEKIASLVLTGQIKDHTLEKILG

CDRAVTVRRLAFDCKLSTLDRGGALSDLPSGPSLDYDRVFGANCEVVVGYIPIPVGMVGP

LTLNDETFFVPMSTTEGCLVASTNRGCKAISQGTGAISTVIRDGITRAPCVRMKSAKEAA

DLKIWCEQKDNFALLKKEFESTTSFGKLLSVSPIVAGKNVYIRLQCFSGDAMGMNMVSKG

SLAVIDLLKSTFPTLELVALSGNVCCDKKSGAINWIEGRGKSVVVEATIPLEVVRSTLKT

NVKTIVHVNIQKNLIGSAMASTVGGFNAHASNIVTAIFLATGQDPAQNVESSNCITLMEE

TDEGDLWISCTMPSIEVGTVGGGTSLPAQSACLKAIGCYGSGDTPGQNAKKLASVVAATT

MAGELSLLAALASNTLVQAHMAHNRKPATSK

>Attheya_septentrionalis

MTNTVASVPTVGERLDALIDTIDSLPSTHLYGAVVAVSVVFSFALLNWGGPSNHTREDAP

QKRRLSWEMRRSAASASAVSFKSGETEPQPKWHILRVLNYAAVGSFAASVVIFCRGASVY

LNDPATLLQFLVGWSLFLCYFFGFFGISFVDPDGMMMSYDPQQSQANNDPQDSTNCKSSS

AKKPLHPPAAATPVCSDGKASCKAPVVAAAALSSSAIKEMSDEQVAQLVVSNQVKDHMLE

KVLDPHRAVAVRRLAFDTKLASLDRGGALDELPSGPSLDYARVFGANCEIVVGYVPLPVG

MVGPLTLNNETVYVPMATTEGCLVASTNRGAKAICQGGGAHSCIVRDGITRAPCVRMKTA

MEAAKLKNWCEEPQHFAKLKAAFESTTSFGKLEQCNPTVAGRNVYLRLRCFSGDAMGMNM

VSKGSLAVIDLLKSEFPTLFLVALSGNMCTDKKPAAINWIEGRGKSVVVEAIIPKDVVRQ

TLKTTVQSICSTNIHKNLIGSAMAGAMGGFNAHAANIVTAIFLATGQDPAQNVESSNCIT

LMEETEEGNLWISCTMPSIEVGTVGGGTSLPAQSACLKAIGCKGGGAKPGQNAQQLAHVV

AAATMAGELSLLAALAANTLVAAHMQHNRKPTTTSK

>Thalassiosira_weissflogii

MAHPSSYERVSSPPPTIGKHIDAVIAKIDSLPSSYVYTLAIAASLTFSFYLLNSGGGSNS

MQMEPHGDVDRASAIKKPKQKSRRSLDEPQPKWHILKLTNYVATSGFLLSVLKFVSNASM

YLNDSTSLLQFLTIWSIFLCYFFGFFGISFIELDDFEQESHPQHQAPVSAINVSHAKHCN

AVKSGSLEPPVKVINVHPHSTSAPVCSDPASFRKPSSTIPGNLKDMSNEEIASLVLDDKI

KDHQLEKLLDPHRAVEVRRLKFDAQLQSLGRGGALAELPHKHDLDYKRVLGANCEIVVGY

VPIPVGMAGPITLNGESVYIPMATTEGCLVASTNRGCKAITQGSGAFSTIVRDGITRAPL

VRLPSAREAAEVLLWIEDENNFKILKEAFESTTSFGKLIEARPTVAGRNVYIRLRCFSGD

AMGMNMISKGSLAVIECLRRKFPQLSLAALSGNMCTDKKAAAINWIEGRGKSVVIEATIP

KEVVRSTLKTSVPAIVEANTNKNLIGSAMAATVGGFNAHAANNVTAVFLATGQDPAQNVE

SSNCITLMEETSDGDLWISCTMPSIEVGTVGGGTGLPAQSACLKAIGVKGGGDSPGDNSK

KLAHVVAAATMAGELSLMAALASNSLVAAHMAHNRKPVSK

>Odontella_Sinensis

MASSSVAYMTVGQRLDSFIESIDPSLSPTTKLYACVVCASVGFSFLLLNGRGSNVGGPDA

SGGPKSASKNESSSSGPSRRSGGGVSAASASSGGGRREPKWGVLKALNVVAALAFLISVI

RFASDASRHMSDSTSLLKFMSIWSALLCYFFGFFGISFVDAEGFSVAGEETKTTEKTTKK

AAGEEEDDDEDGGPGKKTAPAGSRSIPEKPQHANLHPPAPSAPVCSDPTSRAPITKKALP

PNLQGMPDNEIAALVLSGVVKDHQLEKLLSPARAVPVRRIVFERKLSSLNRGGALDELPH

EPSLDYGRVHGANCEIVVGYVPLPCGVIGPLTLDGETVYVPMATTEGCLVASANRGCKAI

SQGGGARSAVIRDGITRAPCVRMRSAMEAAELKLWCEDPANFARLKSAFESTTSFGKLLG

AHPTVAGRNVYLRLRCFSGDAMGMNMVSKGSLAVIELLRSTFPSLELVALSGNMCTDKKA

AATNWIEGRGKSVVVEATIPKDVVRATLKTTVPAMVNTNLQKNLVGSAMAGALGGFNAHA

ANIVTAIFLATGQDPAQNVESSNCITLLEETPTGDLWISCTMPSVEVGTVGGGTGLPAQA

SCLRAIGCKGGGANPGDNARKLARVVAASVLAGELSLLAALAANTLVQAHMIHNRKKPTA

GK

>Rhizosolenia_setigera

XNNPHFSATMTETIGQRXDALLESIDSVPSLQLYAAIVTSVVFSFVILNTGSHNAGLFQN

NEQQRTISSSQRKGSPTMKSNTANYTNTSFDRTKDNDQNQPQIKWYLLKTLNFAVAIGFV

ISITKFSSDASTYITDSISILKFLCAWSLCLCYFLGXFGMSFVDSEDLCSVPTAPPAGAT

SPVAAVHKKQVSKAASFSRPIHPPAMSTHPVCGTGEISSPTTTASLPDDIQSLSNEVAAS

LVVSNKIKDHTLEKHFGPFRAVXXRRLAFETKLASLGHSGALDDLPSGPSLDYKRVFGAN

CEIVVGYVPIPVGMCGPITLNGESVYIPMATTEGCLVASTNRGCKAISAGSGATSVILRD

GITRAPCLRMKSAKEAADLKLWCEEQENFLLLKXAFESTTSFGKLLSAEPTVSGKNVYXR

LRCFSGDAMGMNMISKGSLAVVEXLRELFPTLSLIALSGNMCTDKKAAATNWIEGRGKSV

VIEATIPKDVVTKILKTDVKSIVSTXLQKNLIGSAMAGALGGFNAHAANNVTAIFLATGQ

DPAQNVESSNCITLLEETEDGDLWICCTMPSIEVGTVGGGTGLPAQSSCLKMIGCKGGGE

TPGDNAKQLAHVVAAGTMAGELSLLAALASNTLVAAHMQHNRKPAQKK

>Coscinodiscus_wailesii

MTHAPANQHTTLGMRIDAVLQEIESISPTVLYSVILVASILFSMILLNTGGGRSTEPPRL

RDDVTKPSRKIQRFSDEPQPKWYILKFCNYAAVIGFAFSIFKFAIDATTYINDSTLLMKF

TVCWSLFMCYFFGFFGISFVDADEIEQRAVAGGFDGREGREVERKGILKGSITRKEHEYP

IHSPPQSIHPVCNPNNADIAKKQLPDKSPSSTRISDIKSLSDVEIASLVTSNKIKDHQLE

KLLDPRRAVDVRRIVVDSKLSSLGRGGALADLPSSPSLDYSRVFGANCEIVVGYVPIPVG

MVGPLTLNGETVYIPMATTEGCLVASTNRGCKAISAGSGAHSSIVRDGITRAPCLRLNST

REAADLMLWCEDESNFKLLKQAFESTTSFGKLLRVDPTIAGRNVYLRLKCFSGDAMGMNM

VSKGSLAVVDYLRTVFPTITLVALSGNVCTDKKAAAINWIEGRGKSVVVEATIPQHVVRS

TLKTTVEAIVSTNIQKNLIGSAMAGTVGGFNAHASNIVTAVFIATGQDPAQNVESSNCMT

LMEETPTGDLWISCTMPSIEVGTVGGGTGLPAQAACLKAMGCQGGAADKPGANAKQLAHV

VASATMAGELSLMAALAANTLVQAHMQHNRKPTIGKKA

>Skeletonema_marinoi

MTFSEPTTIGMRIDAVIASLESLPHTNPEILYASLIGASLMFSFLVLNSGGSSMALPDGG

DADSLNKPKKISNAKPVSSSDEPQPKWHILRLTNYVVTFGFLLSVLKFASNATTYLDDST

SLLQFLIIWSISMCYFFGFFGISFIELDELESEPQNHQHQMLRKPMPTEESAEPPVKVVK

MHPPASSAPVCTDPTSFKKPSPSIPSNIKELSNGEIASLVLQDKVKDHQLEKLLDPHRAV

EVRRLKVDNQLDSLGRGGALAELPHKHDLDYKRVLGANCEIVVGYIPIPVGMAGPITLNG

ESVYIPMATTEGCLVASTNRGCKAITQGSGAVSSILRDGITRAPCVRLPSAKEASEVFLW

IERPENFQKLKEVFESTTSFGKLLEARPSVAGKNVYIRLRCFAGDAMGMNMISKGSLAVI

ECLRQVFPNISLVALSGNMCTDKKAAAINWIEGRGKSVVIEATIPKGVVRSTLKTSVPAI

VEANVNKNLIGSAMAGVVGGFNAHASNNVTAVFLATGQDPAQNVESANCITLMEETPEGD

LWISCTMPSIEVGTVGGGTGLPAQAACLKAIGVKGGGENPGDNAKQLAHVVAAATMAGEL

SLMAALASNSLVAAHMTHNRKPASK

>Skeletonema_menzelii

MTSEATTIGMRIDAVIASLESLPHTNPEILYASVIGASLLFSFLVLNSGGSSMALPDGGD

ADSLNKPKISKAKLVSPSDEPQPKWHILRLTNYIVTFGFLLSVLKFASNATTYLDDSTSL

LQFVIIWSISMCYFFGFFGISFIELDELENEPQNHQHQILRKPMPTKESADPPVKVVKMH

PPASSAPVCIDPTSFKKPSPSIPSNIKELSNSEIASLVLQDKVKDHQLEKLLDPHRAVEV

RRLKVDVQLDSLGRGGALAELPHKHDLDYKRVLGANCEIVVGYVPIPVGMAGPITLNGES

IYIPMATTEGCLVASTNRGCKAISQGSGAVSSILRDGITRAPCVRLPSAKEASEVHLWIE

SPENFQKLKEVFESTTSFGKLLEARPSVAGKNVYIRLRCFAGDAMGMNMISKGSLAVIEC

LRQRFPNLSLVALSGNMCTDKKAAAINWIEGRGKSVVIEATIPKGVVRSTLKTSVPAIVE

ANLNKNLIGSAMAGVVGGFNAHASNNVTAVFLATGQDPAQNVESANCITLMEETPEGDLW

ISCTMPSIEVGTVGGGTGLPAQAACLKAIGVKGGGENPGDNAKQLAHVVAAATMAGELSL

MAALASNSLVAAHMTHNRKPASK

>Thalassiosira_miniscula

MPSSPTAPPPPPTLGMQIDAVIASLESISTPQLYGTVIAISLAFSFYLLNSGGGSSSPAL

QMMTNDTKEQQDADADSLRKPARAAATSSSSSSESSSEDRPEPRWHILRFTNYVAALGFL

VSVLQFASNASAYLNDSNSLLQFLTVWSLFVVYFVGFFGISFVELDDFVDGAQISSSVAR

QRAQQQQQQQQQQQSQNMTVAASQAKTPGRKSHKTDGSDSPPIKVVNVHPPATSAPVCSD

PKSFKKPSAASIPPNLKELPNSEIASLVLQDKIKDHQLEKLLDPHRAVQVRRLKFDARLS

ALGNGGALDELPHRHDLEYKRVLGANCEIVVGYVPIPVGMAGPVTLNGESVYIPMATTEG

CLVASTNRGCKAISQGSGASSTILKDGITRAPCVRLPSAKEAAEVALWIETPENFATLKE

AFESTTSFGKLLSARPTVAGKNVYVRLRCFSGDAMGMNMISKGSLAVIECLRQIFPALSL

VALSGNMCTDKKAAAMNWIEGRGKSVVIEATIPADVVRSTLKTSVTAIVEANLNKNLIGS

AMAGTVGGFNAHAANNVTAVFLATGQDPAQNVESSNCITLMEETPEGDLWMSCTMPSIEV

GTVGGGTGLPAQAACLRAIGAKGGGENPGDNARRLAHVVAAATMAGELSLMAALASNSLV

AAHMAHNRKPASK

>Thalassiosira_punctigera

XDTSELKGSKSSNRSPIXPSQNSESMGDATSAPPTVGMQIDAAIAQLENVSTTQLYAIVI

AASLAFSFYLLNSGGGSSALTIMNEKDGDADSLRKPEARKRKDRPRASTSSDEPEPRWHI

LRITNYVVAAGFLLSVLQFASNASTYLNDSTSLLQFLSVWSIFLCYFFGFFGISFIELDD

FVDQQPSQQQQQPSHKLKKAESISPDPPVKVVNVHPPSSSAPVCTDPTSFKKPPSNVPGN

LKDLPNSEIASFVLQDKIKDHQLEKLLDPHRAVEVRRLKFDAKLDSLGRGGALADLPHKH

DLDYKRVLGANCEIVVGYLPIPVGLAGPITLNGESVYIPMATTEGCLVASTNRGCKAISQ

GSGASSTILKDGITRAPCVRLPSAKEAAKVALWIGTPENFLTLKKAFESTTSFGKLLDAT

PTVAGRNVYIRIRCFSGDAMGMNMISKGSLAVIETLKKVFPDLSLLALSGNMCTDKKAAA

LNWIEGRGKSVVIEATIPKDVVRSTLKTSVQAIVEANVNKNLIGSAMAGTVGGFNAHAAN

NVTAVFIATGQDPAQNVESSNCITLMEETSEGDLWISCTMPSIEVGTVGGGTGLPAQSAC

LKVIGVKGGGENPGDNARQLAHVVAAATMAGELSLMAALASNSLVAAHMSHNRKPTSK

>Ditylum_brightwellii

KMATRLTETTDVTIGMRLDELISSIDTIPSTQLYMAVVVVSVIFSFLLLNSGGTSIEQEA

PPSSPPSTALKKITRQGPEPKWHILKILNYAAVSSFLVSVGCFASDASRHMNDSSSLFKF

LFGWSFFLCYFFGFFGISFVDADEIMEGQNFESSERSTSSKSKAVHPPAESTPVCTVPSS

FEKPSLSKTDLQSKSNEELSNMVLTGQIKDHQLEKLLDHHRAVDVRRLAFDSKLSSVGCG

GALTDLPSGPSLDYSRVFGANCEIVVGYVPLPVGMVGPLTLNGQSVYIPMATTEGCLVAS

TNRGCKAITQGSGATSVILRDGITRAPCVRMNSAKEAAELKLWCEMPANFATLKHAFEST

TGFGKLLSVEPTVAGKNAYLRLQCFAGDAMGMNMVSKGSLAVIDLLKSVFPTLVLVALSG

NMCTDKKAAATNWIHGRGKSVVVEAIIPQHVVRTTLKTTVQAIVQTNIHKNLIGSAMAGA

IGGFNAHAANNVTAVFLATGQDPAQNVESSNCITLLEETEDGDLWICCTMPSIEVGTVGG

GTSLPAQSACLKAIGCKGGGVNPGDNAKQLAHVVAAATMAGELSLLAALAANTLVQAHMT

HNRKPTTAKST

>Cylindrotheca_closterium

MASVISNYIDELSSTQLYTAIVGATVTLCVVLLGPAAGGNENLPNNMLGTTTTSNDNNKK

NAQQKQQPKWYLFKYINVGVFCMFVSSVAIFLWDASTYIHNGDRMTQFLVGWSLCLCYFF

GFFGVFFIHQDLLMNQNQNQQGDSSDIAENKQEMVPKTKEVKSVSPPPKAKVVHEPAACA

PVCSDPASFNSSSSSNKKAAISKDTDITGMTNDEIAQLVLDLQVKDHELEKRLDPFRAVT

VRRMIAQHKLQSVHSASSSAFLEDLPEGPSLDYSKVFGTNCEMVVGYVPLPVGMVGPLTL

NGESVYFPMATTEGCLVASTQRGAKAISQGTLGAQALIVKDGITRAPCVRMASAMEAAQL

KLWCQEAANLQVLKEAFESTTSFGKLLECHATVAGKNVYLRLVCFSGDAMGMNMVSKGSL

KVIETLQNIFPNLELVALSGNMCTDKKAAATNWLHGRGKSVVVEAVIPKDVVKKTLKTTV

PALVSTNQNKNLIGSAMAGAMGGFNAHASNIVTAIFLATGQDPAQNVESANCITLMEQVE

ETGDLWISCTMPSIEVGTVGGGTTLPAQAACLEILGCKGGSRENPGKNAQKLALVVAAAT

MAGELSLMSALAANTLVQAHMKHNRKKTV

>Nitzschia_punctata

MAATFDTMPTTIGMKLDAMIAEIDNLSSTQLYGVIVAATVVLCVVLLGTGHSNLDLQHSN

NNDPLLKKQPAVAPSGNIKQPRWHIFKWINYLAVAAFLWSVCTFCLNASQYLHHEESQGV

LVKFLLGWSVFLLYFFGFFGVSLIHEDIPKEEAGAASSAANRLSSVSQSNKTKTVETSSK

NKALHGAAACTPVCSDPSSFKVSSSVPSNIKELADEDVADLVLKNKVKDHELEKRLDPFR

AVTVRRMTANRKLASVLPQNKPNVLDKLPATPSLDYSKVHGANCEIVVGYVPLPVGLVGP

LSLNNETVYVPMATTEGCLVASTNRGAKAITQGGGAQARIVRDGITRAPCVRMESAMGAA

DLKVWCEKPENFARLKQAFEGTTSFGKLQACHPTVAGKNVYLRLVCFSGDAMGMNMVSKG

SLAVIETLQKEFPSLQLVALSGNMCTDKKAAATNWLQGRGKSVVVEATIPKDVVRTTLKT

TVAALVHTNMHKNLIGSAMAGSLGGFNAHASNIVTAVFLATGQDPAQNVESSNCITLMEE

TDEGDLWISCTMPSIEVGTVGGGTSLEAQAACLEAIGCKGGGATPGENAKKLATVVAAAT

MAGELSLLAALAANTLVQAHMTHNRKSNKK

>Nitzschia_sp.

VLSQVDELSSTQLYGLIVAATVTLCVFLLGTGSNTDVEFAATNSKMKTGTNDDGLKKPNA

STNNTASTTRQPRWHIFKWVNYLASAAFLSSVGMFCMNASQYLHHESKGVLVQFLVGWSV

FLMYFFGFFGISLIHEDIPKDNQDDQGIEQPRNTTTRYAAGSSSTTAAAKKQALHTAAPS

APVCSDPASFKASSSSSVPSNLKELPDEDIANLVLMNKLKDHELEKRLDPFRAVTVRRLV

VNQKLSTLLTDGKTNQPVTNVLEKLPSTPSLDYSRVFGANCEIVVGYVPLPVGLVGPLSL

NDETVYVPMATTEGCLVASTNRGAKAITQGGGAQARIVRDGITRAPCVRMASAMEAADLK

VWCEEPQNFAVLKQAFESTTSFGKLQACNPTVAGKNVYLRLVCFSGDAMGMNMVSKGSLA

VIETLQAKFPSLQLVALSGNMCTDKKAAATNWLHGRGKSVVVEAIIPKDVXPWNSQNNSL

SLNDETVYVPMATTEGCLVASTNRGAKAITQGGGAQARIVRDGITRAPCVRMASAMEAAD

LKVWCEEPQNFALLKQAFESTTSFGKLQACNPTVAGKNVYLRLVCFSGDAMGMNMVSKGS

LAVIETLQAKFPSLQLVALSGNMCTDKKAAATNWLHGRGKSVVVEAIIPKDVVRGTLKTT

VDALIFTNTNKNLIGSAMAGSIGGFNAHASNIVTAVFLATGQDPAQNVESSNCITLMEPT

DDGDLWISCTMPSIEVGTVGGGTSLPAQSACLEAIGCKGGGATAGENAQKLARVVAAATM

AGELSLLAALAANTLVQAHMQHNRKPAPKKELTLTPSFVQNNTPIVPFLRKAHLERTYRS

VLGCSQIVIRKSRAPCVRMASAMEAADLKVWCEEPQNFAVLKQAFESTTSFGKLQACNPT

VAGKNVYLRLVCFSGDAMGMNMVSKGSLAVIETLQAKFPSLQLVALSGNMCTDKKAAATN

WLHGRGKSVVVEAIIPKDVVRGTLKTTVDALVHTNTNKNLIGSAMAGSIGGFNAHASNIV

TAVFLATGQDPAQNVESSNCITLMEPTDDGDLWISCTMPSIEVGTVGGGTSLPAQSACLE

AIGCKGGGATPGENAQKLATVVAAATMAGELSLLAALAANTLVQAHMQHNRKPAPKK

>Pseudonitzschia_pungens

MAQTMGTMATIGTRIDNVIAQFDQLPSTQLYGLIVAATVGLCVVLLGTGNSNFELHHQRQ

KQLQQNNDDLKKPKMAAMPSGRQPRWHIFKWINFVAVGAFLCSVFIFCSNASRYLHHESQ

GVLVQFLVGWSVFLMYFFGFFGVSLIHDDIPREEEESLPSKPQTKSVPTNTSAPKKTVHP

PAACTPVCSDPSSFKSSKPSVPENLKELSDDEIAALVLGNKVKDHMLEKLLDPFRAVTVR

RIACNQKLASVLGREDKSNVLEKLPSEPSLDYSRVFGANCEIVVGYVPLPVGLVGPLTIN

DESVYVPMATTEGCLVASSNRGAKAITQGGGARAKIVRDGITRAPCLRMNTAMEAADLKI

WCEKPQNFAILKQAFESTTSFGKLKECNPTVAGKNVYLRLVCFSGDAMGMNMVSKGSLAV

IETLQKYFPTCQLVALSGNMCTDKKAAATNWLYGRGKSVVVECVIPKEVVRTTLKTTVSA

LVHTNLNKNLIGSAMAGAIGGFNAHASNIVTAVFLATGQDPAQNVESSNCITLMEEEVNG

DLWMCCTMPSIEVGTVGGGTSLPAQAACLEAIGCKGGGVTPGANAKKLATVVAAATMAGE

LSLLAALAANTLVQAHMAHNRKPASKK

>Saccharomyces_cerevisiae

MPPLFKGLKQMAKPIAYVSRFSAKRPIHIILFSLIISAFAYLSVIQYYFNGWQLDSNSVF

ETAPNKDSNTLFQECSHYYRDSSLDGWVSITAHEASELPAPHHYYLLNLNFNSPNETDSI

PELANTVFEKDNTKYILQEDLSVSKEISSTDGTKWRLRSDRKSLFDVKTLAYSLYDVFSE

NVTQADPFDVLIMVTAYLMMFYTIFGLFNDMRKTGSNFWLSASTVVNSASSLFLALYVTQ

CILGKEVSALTLFEGLPFIVVVVGFKHKIKIAQYALEKFERVGLSKRITTDEIVFESVSE

EGGRLIQDHLLCIFAFIGCSMYAHQLKTLTNFCILSAFILIFELILTPTFYSAILALRLE

MNVIHRSTIIKQTLEEDGVVPSTARIISKAEKKSVSSFLNLSVVVIIMKLSVILLFVFIN

FYNFGANWVNDAFNSLYFDKERVSLPDFITSNASENFKEQAIVSVTPLLYYKPIKSYQRI

EDMVLLLLRNVSVAIRDRFVSKLVLSALVCSAVINVYLLNAARIHTSYTADQLVKTEVTK

KSFTAPVQKASTPVLTNKTVISGSKVKSLSSAQSSSSGPSSSSEEDDSRDIESLDKKIRP

LEELEALLSSGNTKQLKNKEVAALVIHGKLPLYALEKKLGDTTRAVAVRRKALSILAEAP

VLASDRLPYKNYDYDRVFGACCENVIGYMPLPVGVIGPLVIDGTSYHIPMATTEGCLVAS

AMRGCKAINAGGGATTVLTKDGMTRGPVVRFPTLKRSGACKIWLDSEEGQNAIKKAFNST

SRFARLQHIQTCLAGDLLFMRFRTTTGDAMGMNMISKGVEYSLKQMVEEYGWEDMEVVSV

SGNYCTDKKPAAINWIEGRGKSVVAEATIPGDVVRKVLKSDVSALVELNIAKNLVGSAMA

GSVGGFNAHAANLVTAVFLALGQDPAQNVESSNCITLMKEVDGDLRISVSMPSIEVGTIG

GGTVLEPQGAMLDLLGVRGPHATAPGTNARQLARIVACAVLAGELSLCAALAAGHLVQSH

MTHNRKPAEPTKPNNLDATDINRLKDGSVTCIKS

>Arabidopsis_thaliana

MDLRRRPPKPPVTNNNNSNGSFRSYQPRTSDDDHRRRATTIAPPPKASDALPLPLYLTNA

VFFTLFFSVAYYLLHRWRDKIRYNTPLHVVTITELGAIIALIASFIYLLGFFGIDFVQSF

ISRASGDAWDLADTIDDDDHRLVTCSPPTPIVSVAKLPNPEPIVTESLPEEDEEIVKSVI

DGVIPSYSLESRLGDCKRAASIRREALQRVTGRSIEGLPLDGFDYESILGQCCEMPVGYI

QIPVGIAGPLLLDGYEYSVPMATTEGCLVASTNRGCKAMFISGGATSTVLKDGMTRAPVV

RFASARRASELKFFLENPENFDTLAVVFNRSSRFARLQSVKCTIAGKNAYVRFCCSTGDA

MGMNMVSKGVQNVLEYLTDDFPDMDVIGISGNFCSDKKPAAVNWIEGRGKSVVCEAVIRG

EIVNKVLKTSVAALVELNMLKNLAGSAVAGSLGGFNAHASNIVSAVFIATGQDPAQNVES

SQCITMMEAINDGKDIHISVTMPSIEVGTVGGGTQLASQSACLNLLGVKGASTESPGMNA

RRLATIVAGAVLAGELSLMSAIAAGQLVRSHMKYNRSSRDISGATTTTTTTT

>Homo_sapiens

MLSRLFRMHGLFVASHPWEVIVGTVTLTICMMSMNMFTGNNKICGWNYECPKFEEDVLSS

DIIILTITRCIAILYIYFQFQNLRQLGSKYILGIAGLFTIFSSFVFSTVVIHFLDKELTG

LNEALPFFLLLIDLSRASTLAKFALSSNSQDEVRENIARGMAILGPTFTLDALVECLVIG

VGTMSGVRQLEIMCCFGCMSVLANYFVFMTFFPACVSLVLELSRESREGRPIWQLSHFAR

VLEEEENKPNPVTQRVKMIMSLGLVLVHAHSRWIADPSPQNSTADTSKVSLGLDENVSKR

IEPSVSLWQFYLSKMISMDIEQVITLSLALLLAVKYIFFEQTETESTLSLKNPITSPVVT

QKKVPDNCCRREPMLVRNNQKCDSVEEETGINRERKVEVIKPLVAETDTPNRATFVVGNS

SLLDTSSVLVTQEPEIELPREPRPNEECLQILGNAEKGAKFLSDAEIIQLVNAKHIPAYK

LETLMETHERGVSIRRQLLSKKLSEPSSLQYLPYRDYNYSLVMGACCENVIGYMPIPVGV

AGPLCLDEKEFQVPMATTEGCLVASTNRGCRAIGLGGGASSRVLADGMTRGPVVRLPRAC

DSAEVKAWLETSEGFAVIKEAFDSTSRFARLQKLHTSIAGRNLYIRFQSRSGDAMGMNMI

SKGTEKALSKLHEYFPEMQILAVSGNYCTDKKPAAINWIEGRGKSVVCEAVIPAKVVREV

LKTTTEAMIEVNINKNLVGSAMAGSIGGYNAHAANIVTAIYIACGQDAAQNVGSSNCITL

MEASGPTNEDLYISCTMPSIEIGTVGGGTNLLPQQACLQMLGVQGACKDNPGENARQLAR

IVCGTVMAGELSLMAALAAGHLVKSHMIHNRSKINLQDLQGACTKKTA

>Oryza_sativa

MDVRRGGGGGRIVGAARRALTWGALPLPMRITNGLAMVSLVLSSCDLLRLCSDRERPLGG

REFATVVYLVSLFAHPDAPATTTGDDDDGQGGSRRARPAAAEPAPMHGHGGGMMEADDEE

IVAAVASGALPSHRLESRLGDCRRAARLRREALRRVTGRGVEGLPFDGMDYQAILGQCCE

MPVGYVQLPVGVAGPLLLDGREYHVPMATTEGCLVASVNRGCRAISASGGAFSVLLRDAM

SRAPAVKLPSAMRAAELKAFAEAPANFELLAAVFNRSSRFGRLQDIRCALAGRNLYMRFS

CITGDAMGMNMVSKGVENVLGYLQNVFPDMDVISVSGNYCSDKKPTAVNWIEGRGKSVVC

EAIIKGDVVQKVLKTTVEKLVELNIIKNLAGSAVAGALGGFNAHASNIVTALFIATGQDP

AQNVESSQCITMLEEVNDGDDLHISVTMPSIEVGTIGGGTCLASQAACLNLLGVKGSNHG

SPGANAKRLATIVAGSVLAGELSLLAALASGHLVKSHMMYNRSSKDVAKAAS

>Solanum_lycopersicum

MDVRRRSEEPVYPSKVFAADEKPLKPHKKQQQQQEDKNTLLIDASDALPLPLYLTTNGLF

FTMFFSVMYFLLSRWREKIRNSTPLHVVTLSELGAIVSLIASVIYLLGFFGIGFVQTFVS

RGNNDSWDENDEEFLLKEDSRCGPATTLGCAVPAPPARQIAPMAPPQPSMSMVEKPAPLI

TSASSGEDEEIIKSVVQGKIPSYSLESKLGDCKRAASIRKEVMQRITGKSLEGLPLEGFN

YESILGQCCEMPIGYVQIPVGIAGPLLLNGKEFSVPMATTEGCLVASTNRGCKAIYASGG

ATCILLRDGMTRAPCVRFGTAKRAAELKFFVEDPIKFESLANVFNQSSRFARLQRIQCAI

AGKNLYMRLCCSTGDAMGMNMVSKGVQNVLDYLQNEYPDMDVIGISGNFCSDKKPAAVNW

IEGRGKSVVCEAIITEEVVKKVLKTEVAALVELNMLKNLTGSAMAGALGGFNAHASNIVS

AVFIATGQDPAQNIESSHCITMMEAVNDGKDLHISVTMPSIEVGTVGGGTQLASQSACLN

LLGVKGANREAPGSNARLLATVVAGSVLAGELSLMSAISSGQLVNSHMKYNRSTKDVTKA

SS

>Thalassiosira_rotula

MVVATSAPPTVGMQIDAAIASLETLSTTQLXATVIAVSLXFSFYLLNSGGESSALTMMNE

SSSDSLRKPKXDRKRMKTQQQQSREETDEPEPRWHILRITNYVVATGFTLSVLQFASNAS

TYLNDSTSLLQFLMVWSLFLCYFFGFFGISFIELDDFVDQQQPSQQQPSQQTQVAPSRQL

KKAESSDPPVKVVNIHPPASSAPVCTDPTSFKKPSSKIPSNLKELPNSEIASLVLQDKIK

DHQLEKLLDPHRAVEVRRLKFDAKLDSLGCGGALTELPHKHALDYKRVLGANCEIVVGYI

PIPVGMAGPITLNGESVYIPMATTEGCLVASTNRGCKAISQGSGASSTILKDGITRAPCV

RLPSAREAAEVALWIGKTXNFLSLKEAFESTTSFGKLLDATPTVAGKNVYIRLRCFSGDA

MGMNMISKGSLAVIDALRQIFPRLSLLALSGNMCTDKKAAAMNWIEGRGKSVVIEATIPQ

DVVRSTLKTSVRAITEANVNKNLIGSAMAGTVGGFNAHAANNVTAVFLATGQDPAQNVES

SNCITLMEETPEGDLWISCTMPSIEVGTVGGGTGLPAQSACLKAIGVKGGGENPGDNAKQ

LAHVVAAATMAGELSLMAALASNSLVAAHMAHNRKPASK

>Pseudonitzschia_delicatissima

CKRTFPSXRSNTDLHPSNKNMAETMTPMPTIGMQLDAMIAKVDALPSTQLYGLIVAVTVG

LCVVVLGTGNSNLEMEQQQQDLKKPKMVVVDGGKQPNWHLFKMINYIAVAAFLYSVFIFC

SNASQYLHHGSQGVLAQFLVGWSVFLMYFFGFFGVSLIHDDIPSDTDSVPSQPQTKSVAP

KPVHKPAACTPVCSDPSSFKSSKSSVPENIKELGDEEIADLVLSNKVKDHMLEKLLDPFR

AVKVRRIACNQKLNSVLATNSTNVLDKLPSEHDLDYGRVYGANCEIVVGYVPLPVGLVGP

LTINDESFYVPMATTEGCLVASSNRGAKAIVQGGGARARIVRDGITRAPCLRMNTAMEAA

DLKIWCEEPQNFAILKQAFESTTSFGKLQSCNPTVAGKNVYLRLVCFSGDAMGMNMVSKG

SLAVIEKLQEHFPTCQLVALSGNMCTXKKAAATNWLHGRGKSVIVECIIPKEVVRTTLKT

TVRALVHTNINKNLIGSAMAGAIGGFNAHASNIVTAIFLATGQDPAQNVESSNCITLMEE

QDNGDLWMCCTMPSIEVGTVGGGTSLPAQAACLEAIGVKGGGVVPGANAKKLATVVAAAT

MAGELSLLAALAANTLVQAHMAHNRKPAAKK

>Pseudonitzschia_arenysensis

MAETMTPMPTIGMQLDAMIAQIDALPSTQLYGLIVAATVGLCVVVLGTGNSNLEMEQHDL

KKPKMVPVDGGKQPNWQIFKVINYIAVAAFLYSVFMFCSNASKYLHHESQGVLAQFLVGW

SVFLMYFFGFFGVSLIHEDIPSEVDSVPSQPKSVAPTKPVHPPAACAPVCSDPSSFKAKK

SCVPENIKELGDEEIADLVLANKVKDHMLEKLLDPFRAVTVRRIACNRKLNSVHGNTSNV

LDKLPSEHALDYSRVYGANCEIVVGYVPLPVGLVGPLTINDESFYVPMATTEGCLVASSN

RGAKAICQGGGAKARIVRDGITRAPCLRMNSAMEAADLKIWCEKPANFAILKKAFESTTS

FGKLIECNPTVAGKNVYLRLVCFSGDAMGMNMVSKGSLAVIEKLQEYFPSCQLVALSGNM

CTDKKAAATNWLHGRGKSVVVECIIPKEVVRTTLKTTVAALVHTNVNKNLIGSAMAGAIG

GFNAHASNIVTAIFLATGQDPAQNVESSNCITLMEKQDNGDLWMCCTMPSIEVGTVGGGT

SLPAQAACLEAIGCKGGGATPGANAKQLATVVAAATMAGELSLLAALAANTLVQAHMAHN

RKPAAKK

>Fragilariopsis_kerguelensis

MASTEVMPTLGMRLDAMISHIDRLPSSQLYGVIVAATVGLCVVLLGTGNSNLELQQQQRQ

RQQQMMNVDDLKKPINNRIANVGGKQPKWHIFKWINYLAVGAFLWSVYTFCSNASQYLHH

ESQGVLVQFLVGWSVFLLYFFGFFGISLIHEDIPKEEDETTTMTVSKNIVSRKPYNASTN

STTTMMKKKNSIHPPASCAPVCSDPTSFKSSGASLSSNSVNLEELADEEIANLVLNNKVK

DHELEKRLDPFRAVTVRRIAFNQKIASVLSDNTNKQNNINNTANVLDKLPSTPSLDYSRV

YGANCEIVVGYVPLPVGLVGPLTINDETVYIPMATTEGCLVASSNRGAKAITQGSGAKAR

IVRDGITRAPCIRMRSAMEAADLKLWCEEPSNFLILKQAFESTTNFGKLKECNPTVAGKN

VYLRLVCFSGDAMGMNMVSKGSLAVIETLQNEFPSLQLVALSGNMCTDKKAAATNWLNGR

GKSVVVEAVIPREVVEKTLKTTVKALVHTNINKNLIGSAMAGAIGGFNAHASNIVTAIFL

ATGQDPAQNVESSNCITLIEETDDNDLWISCTMPSIEVGTVGGGTSLEAQSACLEAIGCK

GGGATPGENARKLATVVAAATMAGELSLLAALAANTLVQAHMVHNRKPASNK

>Leptocylindrus_danicus

MAQTADPMTIGTRLDALIASGAEYLNNASQAQVCATLFVSSVAFSFALLNCGKGAHGSSL

DPTLKPLYKNTMEPIKTKVKSPDGSREPRWYIFKMLNYTAVATFSTSVLHFILYSDVYMN

NDAQMMKLMGAWTLFVLYFFGFFGVSFVDTDDHLEGSSPAEDEISEMTADQASPSAVVAV

AKQMPVKAAAPIHPPAPSHPVCSDKNLTKDALTSSSATPAVKKSSSLSLDLQSMTNEELA

DLVLTNKMKDHQLETKLNPTRAVPIRRLVFEKKLASLGHAKSLDELPYEHSLSYERVFGA

NCEIVVGYVPLPVGMVGPCTLNGESVYIPMATTEGCLVASTNRGCKAITAGGGAVSTLLR

DGITRAPCLRFESAAEAAALALWAQEPHNFAKLKAAFESTTSFGKLLSATPTVAGKNCYL

RLKCFSGDAMGMNMVSKGSLAVVDLLRQHFPTLKLVALSGNMCTDKKPAAINWIEGRGKS

VVVEATIPKDIVRTVLKTTVKAIVDTNLQKNLIGSAMSGSVGGYNAHASNIVTAVFLATG

QDPAQNVESSNCITIMEETDDGDLWISCTMPSIEVGTVGGGTSLPAQAACLKAIGVKGGG

DIPGGNARKLAHVVAVATMAGELSLLAALAANTLVQAHMQHNRKPATPAKK

>Chaetoceros neogracile

METQIPTIGMRIDSLISYFDAIPNTSKAFVIICCLFSFFLLNSSPNPAATLNTSTYPGCD

VTKAKTSNKGSSACNTVSKSSNCESEPQPKWHILKLLNVIAVIGLLTSFFWFASNASTYI

NDSNALLKFMALWSGFLCYFFGFFGISFIDVEELERGSSEEEEKQRTNECQKSLFVHKQL

PVHPPASCTPVCSDNLKISTDKPSKPTTFTPSSNSTNTNQKTDIKSLSNSEIVSMVLSNK

IKDHQLEKLLDPYRAVIVRRATFDQKLSTLNQGDALNDLPYEHDLEYSRVFGANCETVIG

YVPLPVGMVGPLTLNGQTVYFPMATTEGCLVASTNRGCKAISQGSGASSTITRDGITRAP

CIRMKSAKEAAELKIWCEIPQNFLTLKAAFESTTSFGKLQSAQVTVAGKNAYVRLCCFSG

DAMGMNMVSKGSLAVIDLLKEKFPTLDLLALSGNMCTDKKAAAINWIEGRGKSVVVESII

PKDVVHATLKTDVKKIVTVNINKNLIGSAMAGSIGGFNAHASNIVSAVFLATGQDPAQNV

ESSNCITLMEETEEGDLWVSCTMPSIEVGTVGGGTGLPAQSAGLKIIGCKGGGEQPGANA

KKLAHVVASAVMAGGPCFFF

>Chaetoceros dichaeta

QHQIQNAPNSGMRSSVVNPPPSLHAPARHTPVCTDPPPNPKPSPTTPLPPPESTSDQAWA

DMVLTNQVKDHQLEKILNPHRAVVVRRLIFQSKLGSLATDESCTSPLDELPYQHSLDYTR

VHGANCETVVGYIPLPVGMVGPLTLNNQTVYFPMATTEGCLVASTNRGCKAISQGSGAIS

TIVRDGITRAPCLKMRSAREACELKLWCELPSNFCKLKEAFESTTGFGKLQSATTMIAGR

NAYVRLCCFSGDAMGMNMVSKGTLAVVDLLRSIFPTLELVALSGNVCTDKKPSAINWIEG

RGKSVVVEATIPKDVVHTTLKTEVKTIVRVNIDKNLIGSALAGSIGGFNAHAANIVTALF

LATGQDPAQNVESSNCMTLMEETDNGDLWISCTMPSIEVGTVGGGTGLPAQKACLKVIGC

QGGGVKPGENSRQLAHVVASATMAGELSLMAALSANTLVQAHMQHNRKPATTSAVDSKQ*

>Chaetoceros affinis

MPMEDIQSLSNDEIASLVLTNKIKDHRLEKLTDPDRAVIIRRAVYSHKLATLPTSNSTTS

NSPIDALPHLHSLNYSKVYGANCETVIGYVPLPVGIVGPITLNNQSVYFPMATTEGCLVA

STNRGCKAITEGGGAVSVITRDGITRAPCVRMESAKDAALLKLWCEDTTNQSENFVQLKK

AFESTTSFGKLEAVNVSIAGRNAYIRLRCFSGDAMGMNMVSKGSLAVIDYLKTIFPSLVL

VALSGNVCTDKKAAAMNWIEGRGKSVVVEATIPRDVVVKTLKTTVKAMVSVNINKNLIGS

ALAGVVGGFNAHASNIVSAVFLATGQDPAQNVESSNCMTLMEETDSGDLWISCTMPSIEV

GTVGGGTGLAAQAACLGIVGCRGGGENPGDNAKQLAHVVAAAVMAGGVSLLGA
